# Merging integrated population models and individual-based models to project population dynamics of recolonizing species

**DOI:** 10.1101/2023.03.14.532675

**Authors:** Lisanne S. Petracca, Beth Gardner, Benjamin T. Maletzke, Sarah J. Converse

## Abstract

Recolonizing species exhibit unique population dynamics, namely dispersal to and colonization of new areas, that have important implications for management. A resulting challenge is how to simultaneously model demographic and movement processes so that recolonizing species can be accurately projected over time and space. Integrated population models (IPMs) have proven useful for making inference about population dynamics by integrating multiple data streams related to population states and demographic rates. However, traditional IPMs are not capable of representing complex dispersal and colonization processes, and the data requirements for building spatially explicit IPMs to do so are often prohibitive. Contrastingly, individual-based models (IBMs) have been developed to describe dispersal and colonization processes but do not traditionally integrate an estimation component, a major strength of IPMs. We introduce a framework for spatially explicit projection modeling that answers the challenge of how to project an expanding population using IPM-based parameter estimation while harnessing the movement modeling made possible by an IBM. Our model has two main components: [1] a Bayesian IPM-driven age- and state-structured population model that governs the population state process and estimation of demographic rates, and [2] an IBM-driven spatial model describing the dispersal of individuals and colonization of sites. We applied this model to estimate current and project future dynamics of gray wolves (*Canis lupus*) in Washington State, USA. We used data from 74 telemetered wolves and yearly pup and pack counts to parameterize the model, and then projected statewide dynamics over 50 years. Mean population growth was 1.29 (95% CRI 1.26-1.33) during initial recolonization from 2009-2020 and decreased to 1.03 (IQR 1.00-1.05) in the projection period (2021-2070). Our results suggest that gray wolves have a >99% probability of colonizing the last of Washington State’s three specified recovery regions by 2030, regardless of alternative assumptions about how dispersing wolves select new territories. The spatially explicit modeling framework developed here can be used to project the dynamics of any species for which spatial spread is an important driver of population dynamics.

## INTRODUCTION

Estimates of abundance and vital rates are key outputs of most population models, and are of particular importance in modeling small, threatened populations (Schaub et al., 2007). In certain recovering populations, one of the most consequential vital rates is the rate of colonization of unoccupied habitat. Observing colonization is difficult and yet it is of great interest from a management perspective (e.g., Recio et al., 2020), particularly in cases where recovery goals are spatially explicit (e.g., Maletzke et al. 2016). Modeling the colonization of a landscape over time, which requires accounting for both demographic processes and often complex spatial processes driven by individual behavior and movement, is an underexplored challenge.

Integrated population models (IPMs) are a powerful and increasingly indispensable tool for modeling population dynamics (Schaub & Kery, 2021). IPMs integrate various data streams (e.g., mark-recapture, count, and reproductive data) in a single analytical framework, offering advantages of increased precision and more efficient use of data (Abadi, Gimenez, Arlettaz, et al., 2010; Schaub & Abadi, 2011). IPMs have been applied to a wide variety of taxa to answer questions related to population viability (Saunders et al., 2018), immigration (Abadi, Gimenez, Ullrich, et al., 2010), disease dynamics (Gamble et al., 2020), and more. Traditional IPMs are age-or state-structured and are non-spatial or have limited discrete spatial states. Thus, traditional IPMs are not capable of capturing the complex spatial processes, frequently determined by individual behavior and movements, that drive colonization. While fully spatial IPMs have been described (e.g., Chandler and Clark 2014), the data requirements for building such models are in many cases prohibitive, especially for modeling populations at large spatial scales. Furthermore, the integration of complex behavioral and movement dynamics into demographic estimation models is in its infancy (but see Converse et al. 2022). Thus, existing IPM frameworks are not adequate for populations where colonization is of paramount importance.

While IPMs are not typically structured to accommodate individual processes such as behavior and movement, individual-based models (IBMs) are regularly used to account for such processes in the study of population dynamics (Grimm, 1999). IBMs allow individuals to have particular states (e.g., spatial location, age, reproductive status) and rates (e.g., probabilities of survival, fecundity, and movement) that are updated iteratively and produce emergent predictions at the population level (Grimm, 1999), but do not traditionally integrate an estimation component. Furthermore, they typically require substantial computing resources, especially if a full representation of parametric uncertainty is desired (Grimm, 2019). Thus, there remains an analytically unexplored space in which demographic processes (as estimated by an IPM) and movement processes (modeled using an IBM) could be combined within a single modeling framework to efficiently and effectively project population growth and spread.

To address this gap, we developed a spatially explicit population modeling framework that integrates individual-based movement with an IPM, allowing us to harness the benefits of IPMs while accounting for the individual-based dynamics that drive colonization. Our model combines multiple data streams (e.g., counts, encounter histories of marked individuals) within the IPM component and incorporates finer-scale behavioral and movement processes into the individual-based component. In application to threatened populations, our model represents a population viability analysis (PVA; Morris & Doak (2002)) and can be expanded to model scenarios related to management and structural uncertainties to determine their impacts on future abundance and progress toward recovery goals.

The focal species and motivation for the development of this model was the gray wolf (*Canis lupus*) in Washington, USA, a state-endangered, recovering species that is recolonizing the state. Dispersing wolves can travel hundreds of kilometers searching for potential mates and new territories (Boyd & Pletscher, 1999; Jimenez et al., 2017), allowing the species to recolonize new areas quickly, but also making it difficult to accurately model the recolonization process. The goals of our analysis were to [1] combine data from population counts and Global Positioning System (GPS) collars to estimate current population dynamics of wolves in Washington, and [2] project the population forward within a PVA framework to assess future population trajectories and progress toward statewide recovery. To capture fundamental structural uncertainty about how dispersing wolves select new territories, we developed and compared two different approaches for modeling this process. Our framework can be customized for a wide variety of applications and provide valuable information for management in cases where individual-based dynamics must be accommodated to realistically project population outcomes.

## METHODS

### Study species

Gray wolves are a pack-forming apex predator native to all northern landscapes with an adequate prey base (Mech, 1970). Though extirpated from much of their range outside wilderness areas by the 1950s (Mech, 1995), many wolf populations are increasing due to increased legal protections (Mech, 1995) and associated recolonization (Chapron et al., 2014) and reintroduction (Fritts et al., 1997). Within Washington, USA, wolves were historically common but were extirpated by the 1930s. The first sign of their recolonization in Washington was the establishment of a resident pack in 2008 (WDFW et al., 2021).

As of December 2022, gray wolves were listed as endangered throughout the state by the State of Washington and in the western two-thirds of Washington under the U.S. Endangered Species Act (WDFW et al., 2021). Washington’s Wolf Conservation and Management Plan identified three recovery regions: Eastern Washington, the Northern Cascades, and the Southern Cascades and Northwest Coast (Wiles et al., 2011). As of December 2022, there were wolf packs in the former two but not the latter recovery region (WDFW et al. 2021; Figure 1).

**Figure 1.**
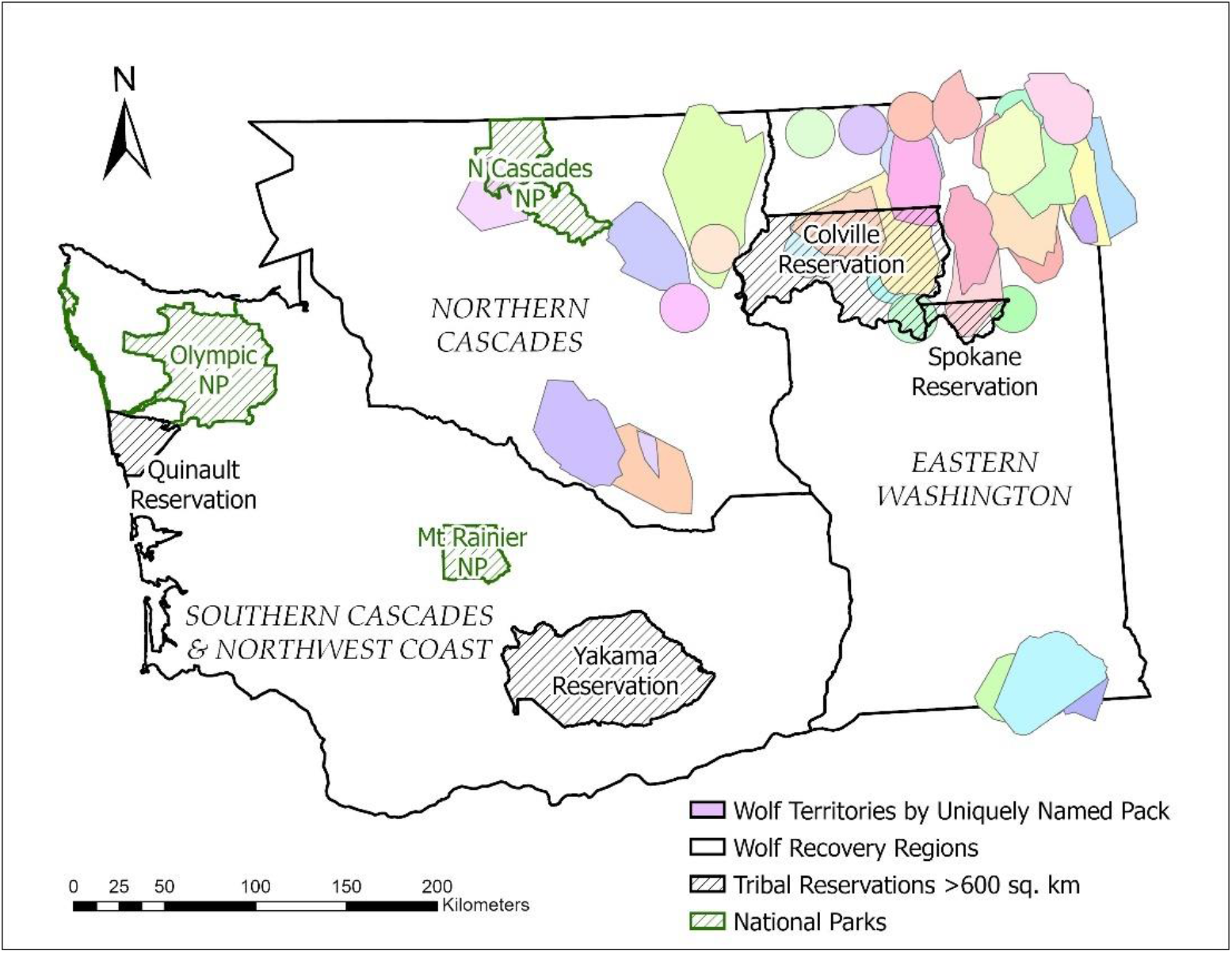
Thirty-eight uniquely named packs in three wolf recovery regions in Washington State during the data collection period (2009-2020). These 38 named packs correspond to 34 pack territories occupied during annual surveys conducted in winter each year. Some pack names changed for the same territory due to the pack naturally dissolving or agency removals due to livestock depredations. Packs represented by circles are those that were known to exist but for which individual movement data were not available.

### Data

Global Positioning System (GPS) collars were placed on 74 individual wolves by the Washington Department of Fish and Wildlife (WDFW) from 2009 to 2020. Nine wolves were re-collared over this duration, for a total of 83 collaring events. Collaring events had a mean duration of 397 (SD = 266, range = 30-1492) days. These wolves represented 38 uniquely named packs in Washington over the period 2009-2020, where a “pack” was defined as two or more wolves traveling together in winter (WDFW et al., 2021). These 38 packs were believed to represent all wolf packs inside Washington’s borders, including on Tribal lands. Wolves were captured in accordance with WDFW’s internal protocols and the guidelines of the American Society of Mammologists for the use of live animals in research (Sikes, 2016).

The minimum known number of wolves in each pack and year (2009-2020) was determined via three methods at the end of each year: aerial surveys, track surveys, and camera surveys. Minimum known number of wolves was calculated as the minimum number observed among the methods used, after accounting for known mortalities that occurred during the survey period. Please see Appendix S1 for more details on wolf captures and surveys.

### Model overview

Our population projection model included two primary processes: demographic processes governed by an IPM and colonization processes governed by an IBM. The general workflow for model development is illustrated in Figure 2. Our first step was using GPS collar data to identify movement states, which allowed us to distinguish, broadly, “movers” and “residents” (see Appendix S2). The identification of movement states allowed us to identify a distribution of dispersal distances for movers along with estimated mean territory size of resident wolves (see Appendix S2). Movement states also provided structure for our Bayesian IPM, which integrated data relevant to estimating survival, state transitions, reproduction, and abundance within an age-and state-structured population model during the data collection period (2009-2020; see *Integrated population model* and Appendix S3). The demographic parameter estimates from the IPM then fed into the projection model.

**Figure 2.**
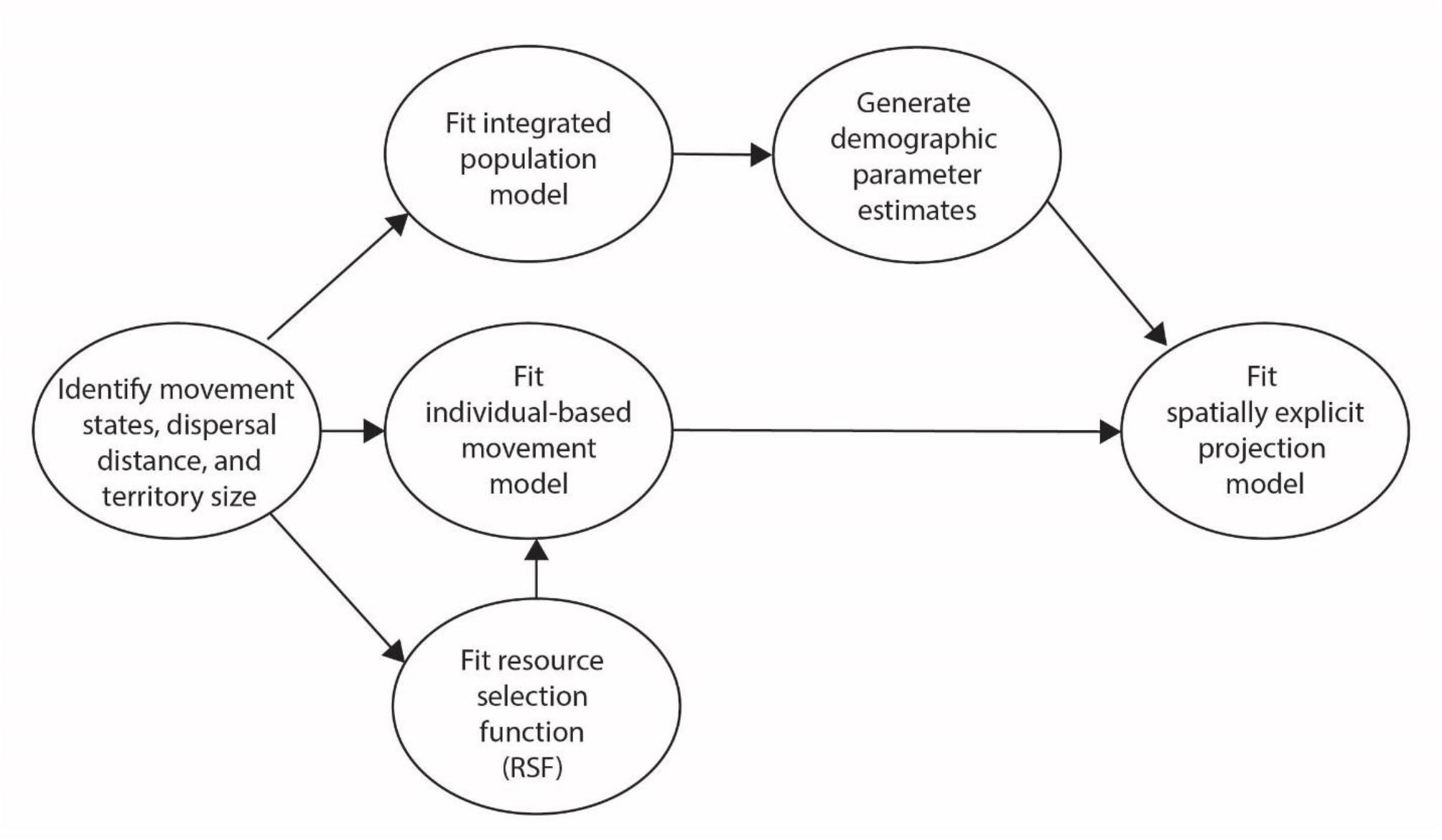
Overview of steps in the construction of our spatially explicit projection model. Identification of movement in GPS-telemetered wolves led to estimation of [1] dispersal distances of movers, [2] mean wolf territory size following removal of movement paths, and [3] state transitions in our multi-state model. The former two components, combined with a resource selection function (RSF) of the second order, informed our individual-based model (IBM), while the latter was a component of our integrated population model (IPM; Figure 3). This IPM included an age and state-structured demographic model which, combined with the IBM, comprised the spatially explicit projection model.

Our IBM incorporated information on dispersal and habitat suitability to model the colonization process across hypothetical territories statewide (see *Individual-based movement model for spatial projection*). We assessed territory-level habitat suitability in Washington State using GPS collar data and a second-order resource selection function (Manly 2002). The IBM was then combined with the demographic parameter estimates from the IPM to model future population dynamics in the projection period (2021-2070; Figure 2; see *Population projections*). In the projection period, individuals were able to colonize new territories in Washington.

### Integrated population model

The IPM component of our projection model is a spatially implicit, age-and state-structured demographic model, fit in a Bayesian framework, including sub-models for [1] survival and state transitions, [2] abundance, [3] reproduction, and [4] the population growth process. We used the GPS collar data to estimate survival and state transitions for “resident” and “mover” wolves in a known fate multi-state model (Brownie et al., 1993; Kaplan & Meier, 1958; Appendix S3). To estimate abundance, we used a log-normal model of the minimum known number of wolves in each territory and year (Appendix S3). Lastly, we estimated fecundity (i.e., number of six-month-old pups in December) as a categorical distribution using the minimum known number of six-month-old pups per wolf pack territory from 2009 to 2014 (Figure 3; Appendix S3).

**Figure 3.**
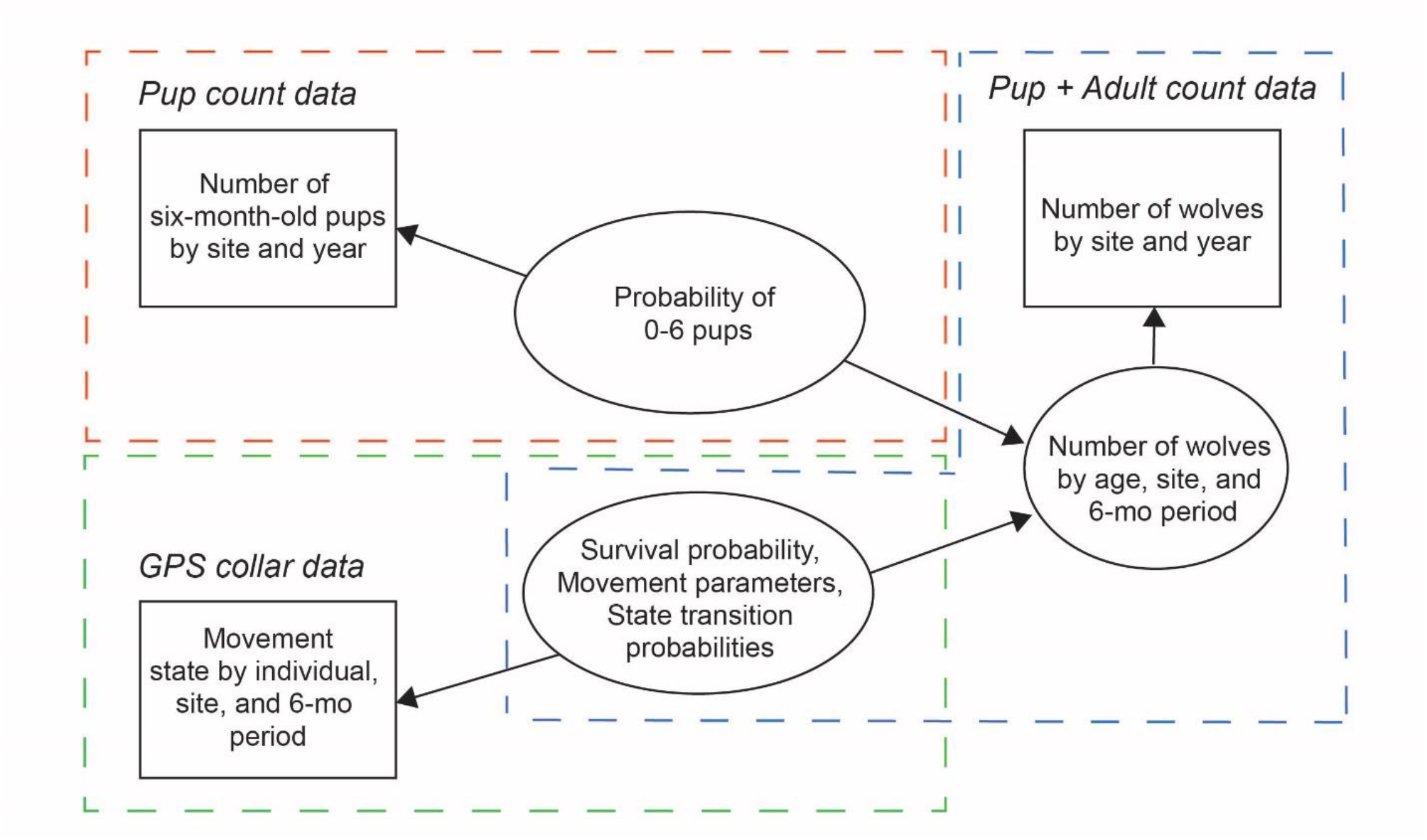
Overview of the integrated population model used to estimate demographic rates and abundance of wolves in Washington State, USA. Winter survey data in December in territory *s* comprised minimum counts of six-month-old pups and minimum counts of all individuals (pups and adults) in each pack. GPS collar data comprised data from individual *i* in period *t* (December or June) and territory *s*. These data streams were used to estimate the probability of a pack having 0-6 pups in December, survival parameters for age grouping *g*, movement state for individual *i* in time period *t*, state transition probabilities (probabilities of transitioning into one of the mover states in the multi-state model for age grouping *g*), probability of staying in Washington, and abundance *N* for age class *age* in time period *t* in territory *s*.

#### Population process model

We used the estimated mean pack territory size to define a spatial surface of potential pack territories across Washington (Appendix S2). Known packs from 2009-2020 were assigned to the nearest potential territory. During the data collection period (2009-2020), wolves were allowed to transition between resident and mover states, but between-territory movements were latent and were only allowed to and from a set of territories known to be occupied.

The model operated on a six-month time step, and we assumed that wolves were born in June. Thus, wolves entered the model in December at six months of age. In December of each year, individuals were age six months (age class 1), 18 months (age class 2), or ≥30 (hereafter 30+) months (age class 3), and in June of each year individuals were age 12 months (age class 1), 24 months (age class 2), or 36+ months (age class 3).

In each territory *s* that was occupied during the data collection period, we modeled age-specific wolf abundance beginning in the year wolves were first counted in that territory. The first sampling period in the population process model was December 2009, and territories could first become occupied in December of subsequent years (i.e., *first*_*s*_ ∈ 1,3,5,, … .23).

Detailed information on the population process model is presented in Appendix S3. Broadly, for *t* > *first*_*s*_, the total number of wolves in six-month period *t* in territory *s* was calculated as the sum of the number of individuals in each age class:

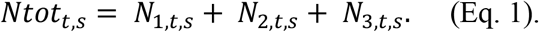

Due to seasonal differences in how data were collected and how animals aged in the model, we had different process models for odd-numbered periods (i.e., December) and even-numbered periods (i.e., June). For *t* = 1,3,5, … .23, the abundance of wolves in the first age class was calculated as:

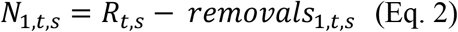

where *R*_*t,s*_ is the estimated number of six-month-old pups in December of each year and *removals*_1,*t,s*_ are animals that were removed by the state agency prior to 6 months of age. For *t = 2,4, 6* … *22*, the abundance of wolves in the first age class was modeled as:

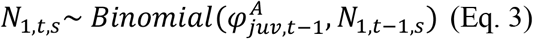

using random binomial draws to account for demographic stochasticity. Note that 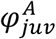 refers to survival of resident juveniles, as survival was estimated separately for juveniles and adults in the multi-state model (see Appendix S3).

For all *t*, abundances for age classes 2 and 3 were calculated as:

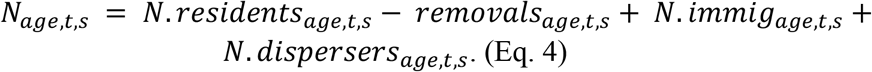

That is, these age classes were composed of residents who survived and remained (*residents*) minus individuals that were removed by the state agency (*removals*) plus immigrants entering Washington from out of state (*immig*) plus dispersers from within Washington who joined the pack (*dispersers*). Estimation of *N. residents*_*age,t,s*_, *N. immig*_*age,t,s*_, and *N. dispersers*_*age,t,s*_ is presented in Appendix S3. Removals for each age class, time period, and territory, *removals*_*age,t,s*_, were provided as data by Washington Department of Fish and Wildlife.

#### Model fitting

For data from 2009-2020, we fit the IPM in a Bayesian analytical framework using JAGS v.4.3.1 (Plummer 2017) through the R package jagsUI (Kellner 2021) in R v.4.2.0 (R Core Team 2022). We ran 3 chains for 240,000 iterations, including 120,000 iterations of burn-in and a thinning rate of 6. We determined convergence by checking all chains visually and calculating the R-hat for each parameter (Gelman & Rubin, 1992). All parameters had R-hats < 1.1, which we took as an indicator of convergence.

### Individual-based movement model

For 2021 and beyond, the movement component was spatially explicit and allowed the colonization of new pack territories in Washington. This was accomplished via an individual-based movement model in which each mover was assigned a dispersal distance and underwent a probabilistic process of settling within a new territory. We note that wolves could join a territory regardless of whether that territory was already occupied.

Given substantial uncertainty in the recolonization process, we used two territory selection processes to assign each mover to a new territory in the forward projections. Common to both methods, each mover was assigned a dispersal distance as a random draw from a gamma distribution. Once the distance was selected, we identified all potential territories at that land-based distance from the origin territory (i.e., dispersing animals were not allowed to cross marine waters, specifically Puget Sound). Both methods relied in different ways on a second-order resource selection function (RSF; Manly et al. 2002). This RSF, based on the daily GPS collar data and with movement trajectories removed, incorporated 11 range-wide covariates hypothesized to influence wolf territory locations (Appendix S4).

In the first approach, termed the “RSF Categorical” method, the destination territory was determined via a draw from a categorical distribution, where the categorical probabilities were calculated from the territory-specific habitat suitability indices predicted from the RSF, standardized to sum to one across territories. The origin territory was included in this process, such that there was a non-zero probability of the wolf returning to its origin.

In the second approach, termed the “Least Cost Path” method, rather than considering destination suitability, animals were assigned to the territory at the selected distance that had the path of least travel cost --that is, we considered the intervening matrix from origin to destination rather than suitability at the destination only. Least-cost paths between each pair of territories were calculated via a least-cost path analysis in program UNICOR (Landguth et al., 2012), with resistance values calculated by inverting the second-order RSF. The probability of returning to the origin was calculated based on the proportion of observed GPS collar trajectories classified as “movers” that terminated in the wolf’s original territory. Thus, each wolf had the potential to colonize a new territory or to return to their source territory.

For both territory selection methods, we incorporated an attraction function to better capture the effect of sociality on pack formation. Once all dispersing wolves were assigned to a territory, if there were single wolves in directly adjacent territories, those two wolves were joined in one of the two territories, with the territory of occupation selected at random.

### Population projections

The projection model used the age and state structure of the population process model, combined with the movement process of the IBM, to estimate wolf population dynamics at future time steps. Every six months, both the population process model and the IBM were engaged to allow for both demographic change and movement. The projection model maintained the annual rate of removals present in the estimation model and maintained the same number of wolves immigrating into Washington from out of state as was estimated across the data period.

Population recovery was defined under Washington’s Wolf Conservation and Management Plan as four breeding pairs (where a breeding pair consists of a minimum of two adults and two six-month old pups alive on December 31^st^) in each of three recovery regions and three additional breeding pairs anywhere in Washington (Wiles et al., 2011). Quasi-extinction, as defined by WDFW, was <46 breeding females in the state and <12 breeding females in each of the three recovery regions (Wiles et al., 2011). We assumed a 1:1 sex ratio, such that quasi-extinction in the model occurred with <92 adult wolves statewide and <24 adult wolves in each recovery region.

The projection model was run in R v.4.2.0 (R Core Team 2022) for 50 years, using 500 samples from the posterior distributions of the model parameters with 100 stochastic simulations run per sample. Metrics for each simulation included total number of wolves, geometric mean of population growth rate, probability of reaching the recovery threshold across all years (2021-2070), probability of dropping below the quasi-extinction threshold across all years (2021-2070), and the probability of extinction, i.e., no wolves at year 50.

## RESULTS

### Wolf Movement

Mean territory size of wolves in Washington was 760.03 (SE = 57.12) km^2^, calculated from territories corresponding to 21 unique packs and 55 pack-years with >180 daily points. This led to the creation of 224 potential territories across Washington. These potential territories were highly variable in their habitat suitability.

The distribution of one-way movement distances (n = 34; median = 105 km, max = 632 km) for Washington wolves was modeled as a gamma distribution with shape and rate of 0.814 and 0.005, respectively. The probability of a mover wolf returning to its origin territory was 5/34 = 0.1471. Twelve to 24-month-olds had a ∼6.5x higher probability of initiating movement in any six-month period compared to ≥24-month-olds, while ≥24-month-olds had a ∼1.3x higher probability of continuing movement compared to 12-to 24-month-olds (Table 1). The probability of staying within Washington in a movement state was 0.75 (95% Credible Interval [CRI] = 0.64 -0.85; Table 1).

**Table 1.** Posterior mean and 95% credible intervals for parameters from the integrated population model for wolves in Washington State, USA, using GPS collar and winter pack and pup count data from 2009-2020.

| Parameter | Description | Mean (95% Credible Interval) |
| --- | --- | --- |
| $\varphi_{juv}^A$ | 6-mo survival of residents 6-24-mo-olds | 0.98 (0.91 - 1.00) |
| $\varphi_{ad}^A$ | 6-mo survival of residents $\geq 24$ mos | 0.94 (0.89 - 0.98) |
| $\varphi_{juv}^B$ | 6-mo survival of movers 12-24 mo-old | 0.84 (0.46 - 1.00) |
| $\varphi_{ad}^B$ | 6-mo survival of movers $\geq 24$ mos | 0.98 (0.92 - 1.00) |
| $\epsilon_{juv}^A$ | Prob of initiating movement for residents 12-24 mos | 0.38 (0.25 - 0.52) |
| $\epsilon_{ad}^A$ | Prob of initiating movement for residents $\geq 24$ mos | 0.06 (0.03 - 0.10) |
| $\epsilon_{juv}^B$ | Prob of continuing movement for movers 12-24 mos | 0.33 (0.26 - 0.42) |
| $\epsilon_{ad}^B$ | Prob of continuing movement for movers $\geq 24$ mos | 0.43 (0.35 - 0.51) |
| $\alpha$ | Prob of mover wolf staying within WA State | 0.75 (0.64 - 0.85) |
| $\pi_1$ | Prob of a territory with 2+ wolves having 0 6-mo-olds | 0.50 (0.38 - 0.62) |
| $\pi_2$ | ...1 6-mo-olds | 0.07 (0.02 - 0.16) |
| $\pi_3$ | ...2 6-mo-olds | 0.14 (0.06 - 0.25) |
| $\pi_4$ | ...3 6-mo-olds | 0.17 (0.09 - 0.26) |
| $\pi_5$ | ...4 6-mo-olds | 0.08 (0.03 - 0.15) |
| $\pi_6$ | ...5 6-mo-olds | 0.02 (0.00 - 0.06) |
| $\pi_7$ | ...6 6-mo-olds | 0.02 (0.00 - 0.05) |
| $\lambda.immig_{data}$ | # of new immigrants / occupied territory in 6-mo period | 0.11 (0.01 - 0.28) |

### Wolf Demography

Average pack size during the data period (2009-2020) was 4.67 ± 2.54 (1 SD), with an average of 1.50 ± 1.73 six-month-old pups per pack. Six-month survival probability for age grouping 1 (6-to 24-month-olds) was 0.98 (95% CRI = 0.91 - 1.00) for state 1 (residents) and 0.84 (0.46 - 1.00) for state 2 (in-state movers; Table 1; but note that six-month-old pups were not permitted to move). Six-month survival probability for age grouping 2 (≥24-month-olds) was 0.94 (0.89 - 0.98) for residents and 0.98 (0.92 - 1.00) for in-state movers (Table 1).

Occupied territories were most likely to produce zero (π_1_ = 0.50; 95% CRI = 0.38 - 0.62), three (π_4_ = 0.17; 0.09 - 0.26), or two (π_3_ = 0.14; 0.06 - 0.25) six-month-old pups in December (Table 1). Rate of immigration, λ. immig_data_, was estimated to be 0.11 (0.01 - 0.28), equating to approximately one new out-of-state immigrant entering each occupied territory every nine years.

We estimated a total of 172 wolves (95% CRI = 155 - 191) in Washington in December 2020, with a geometric mean growth rate (λ. growth) of 1.29 (1.26 - 1.33) from 2009 – 2020 (Figure 4).

**Figure 4.**
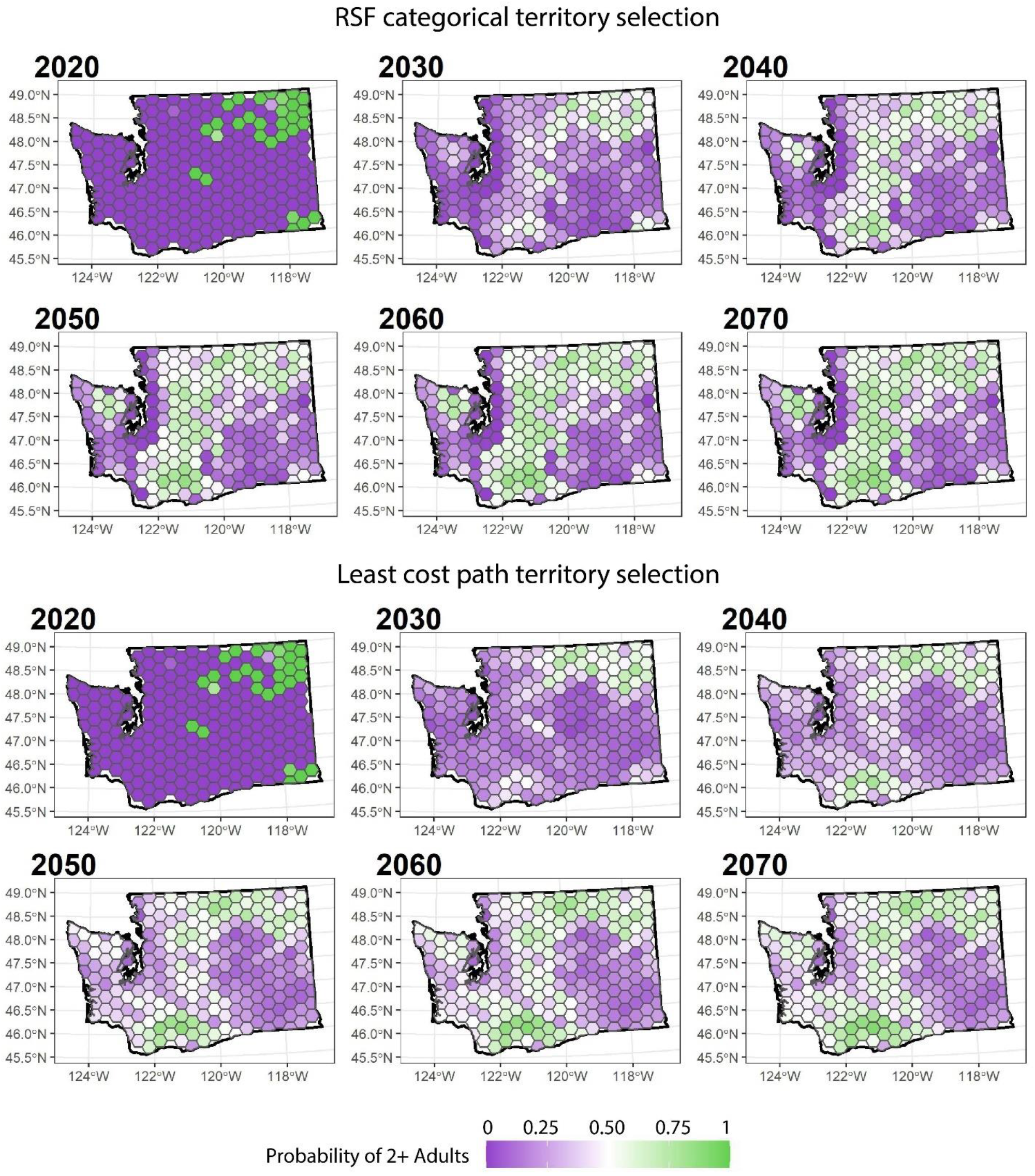
Probability of having ≥2 adults, by wolf territory, for each of two alternative methods for modeling how dispersing wolves select new territories. The resource selection function (RSF) Categorical method considered destination only and included a territory selection process based on median RSF values at a given dispersal distance from the origin territory. The Least Cost Path method considered the intervening matrix and included a territory selection process based on the relative cost of getting to a territory at a given dispersal distance.

### Population Projections

Spatial spread at various time points differed somewhat between the two territory selection methods, RSF Categorical and Least Cost Path (Figure 4). Both methods predicted that the currently uninhabited Southern Cascades and Northwest Coast recovery region would have ≥ 1 territory with 2+ adults by 2030 (RSF: 99.7% of simulations; Least Cost Path: 99.4% of simulations). However, while the RSF Categorical method predicted wolf territories in the Olympic Peninsula by 2040, the Least Cost Path method predicted that this would occur later, by year 2050 (Figure 4).

While spatial spread varied depending on the territory selection method, metrics related to population growth, progress toward recovery, and probabilities of quasi-extinction and extinction were practically identical for both territory selection methods (Appendix S5). Therefore, we focus on results related to the RSF Categorical method only.

Wolf abundance at the state level increased from a median of 271 (95% prediction interval [PI] 77-523) in 2030 to 544 (54-1363) in 2070 (Figure 5), with λ. growthof 1.03 (1.00-1.05) over the projection period (2021-2070).

**Figure 5.**
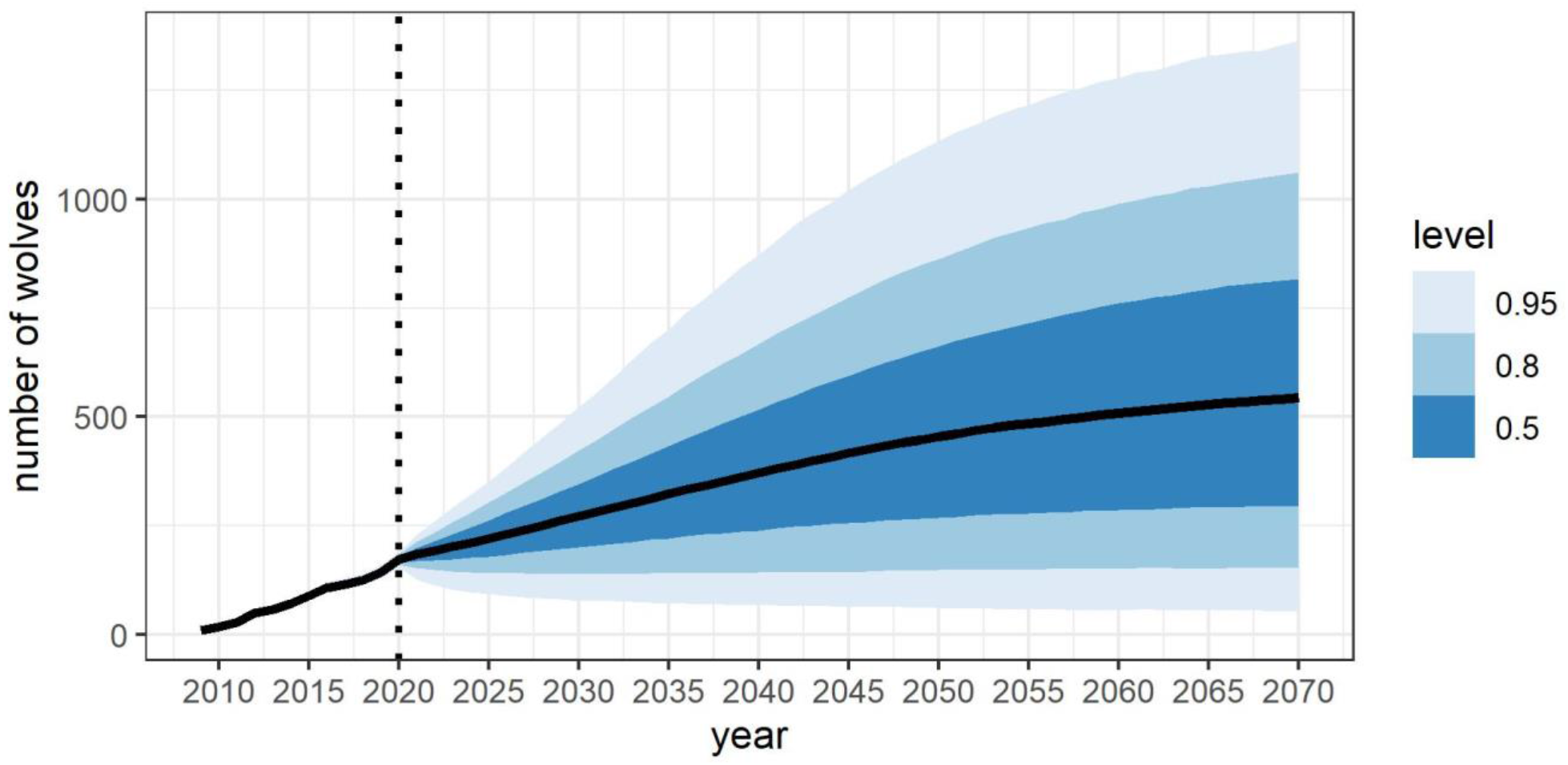
Estimated number of wolves in Washington State, USA, including both the data collection period (2009-2020) and the projection period (2021-2070), separated by dotted line. Black line indicates the mean, and gradations of blue represent abundance estimates within the 50% credible interval (CRI), 80% CRI, and 95% CRI.

Median probability of recovery (i.e., four breeding pairs in each recovery region, with three additional breeding pairs anywhere in the state) across all years (2021-2070) was 0.72 (95% PI 0.00-1.00). This probability of recovery increased over time, from 0.00 (0.00-0.00) in 2020 to 0.94 (0.02-1.00) in 2070 (Figure 6).

**Figure 6.**
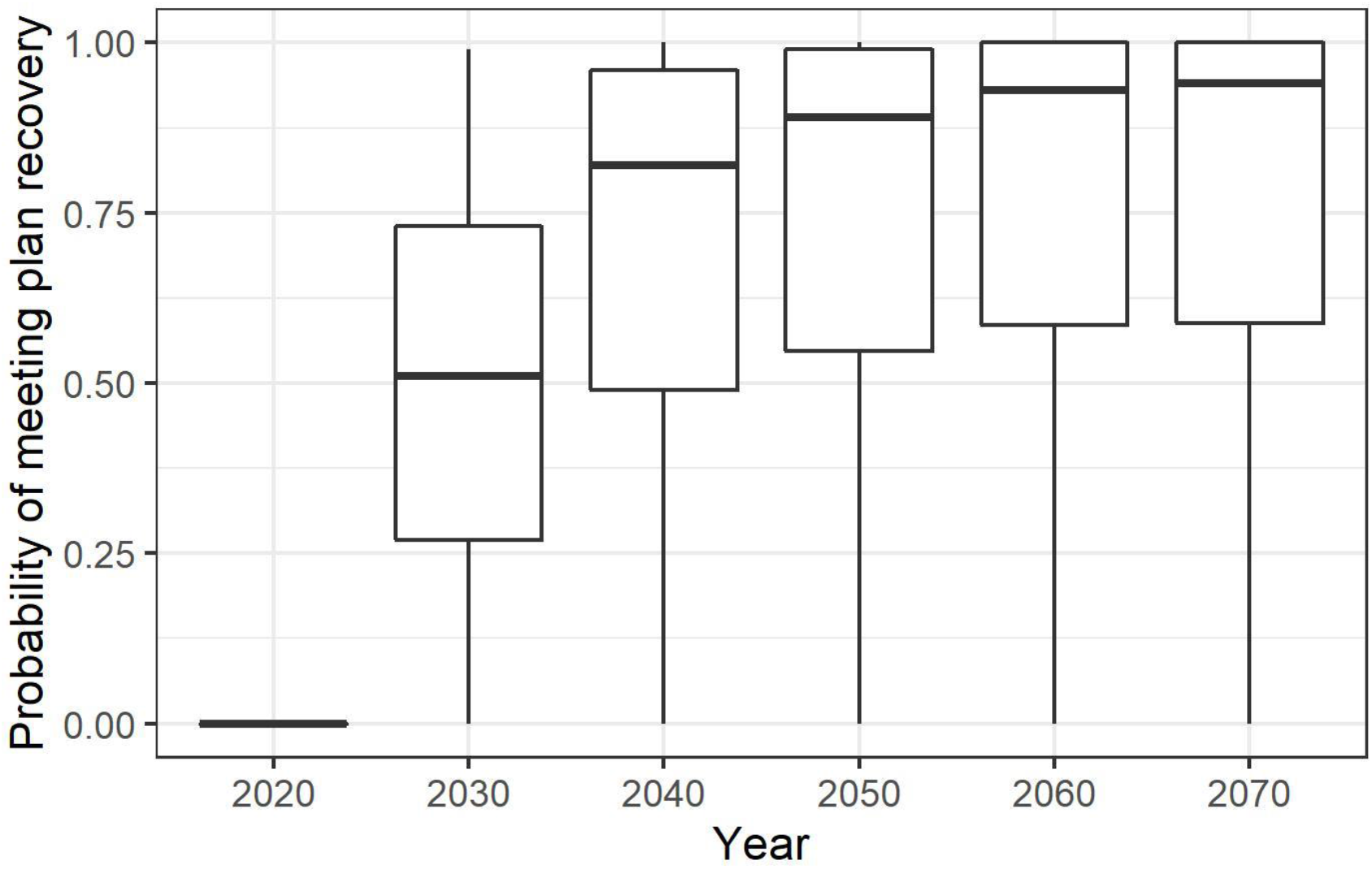
Probability of meeting plan recovery at various time points. Plan recovery is considered having four breeding pairs in each recovery region, with three additional breeding pairs anywhere in the state. The center line represents the median.

Median probability of quasi-extinction across all years (i.e., <92 adult wolves in the state and <24 adult wolves in each recovery region from 2021-2070), as well as median probability of extinction (i.e., zero wolves in 2070), were close to 0 ([0.00, 95% PI 0.00-0.32] and [0.00, 95% PI 0.00-0.00], respectively).

## DISCUSSION

Our spatially explicit model, by combining an age- and state-structured IPM with an IBM for modeling spatial spread, projected the dynamics of recolonizing wolves efficiently. Both the estimation and projection components of the model accounted for uncertainty across the parameters of interest. We developed a model framework that is particularly suited to conducting PVAs and can be easily expanded to evaluate the effects of management strategies or key model uncertainties while estimating future population trajectories and progress toward recovery goals.

We found that while gray wolf populations grew at around 29% from 2009-2020 in Washington, this rate was only around 3% from 2021-2070. A lower growth rate during the projection period was likely driven in part by a decreasing relative number of out-of-state immigrants, as we held the total number of immigrants per period constant as the population grew during the projection period. While we did not have an explicit density-dependent component in our model, there were limited potential territories in suitable habitat, and the spatially explicit dispersal process potentially acted as a regulator on total population size by spreading animals out in space. As a balance to the dispersal process, our attraction function allowed individual animals in adjacent territories to form pairs, reducing the number of lone wolves in territories.

Estimation of latent parameters, particularly out-of-state immigration, was made possible by the integrated estimation framework (Abadi, Gimenez, Ullrich, et al., 2010). Immigration has been a key driver of wolf recolonization of Washington (Maletzke et al., 2016; Wiles et al., 2011), though harvest rates in Idaho, Montana, and British Columbia may alter immigration in the future. Estimating latent immigration and incorporating it into the projection model were essential for understanding wolf population dynamics and recovery potential.

In addition to understanding the gray wolf population trajectory in Washington, an important aspect of our work was predicting the distribution of wolves across the state. Maletzke et al. (2016) predicted the establishment of wolf packs in the Southern Cascades and Northern Coast recovery region by 2021; though this has not yet occurred, their work demonstrated the need to focus on dispersal. While we found that our two methods for modeling territory selection by dispersing wolves had similar effects on population growth, they had differing impacts on spatial spread. In particular, the timing of wolves entering the Olympic Peninsula differed between the two methods: 20 years for the RSF Categorical method and 30 years for the Least Cost Path method. The RSF Categorical method considered only the relative suitability of potential destination territories, while the Least Cost Path method considered the cost of traversing the matrix to a destination territory. Thus, the Least Cost Path method produced more conservative results, making it less likely for wolves to arrive in areas requiring travel across a matrix of poor-quality habitat (e.g., the I-5 freeway corridor running south from Puget Sound). There is evidence that carnivore dispersal movements are influenced by the quality of habitat between source and destination (Elliot et al., 2014; Rio-Maior et al., 2019). However, there is also evidence that dispersing carnivores are capable of traversing long distances in suboptimal landscapes (Jimenez et al., 2017) and with greater tolerance for risk than when in a resident state (Barry et al., 2020). We propagated this uncertainty in how the habitat matrix influenced dispersal into uncertainty in the timing of wolf colonization of parts of Washington.

Explicitly incorporating greater complexity in the model could allow for more realistic population dynamics. Density dependence can affect dispersal (Morales-González et al., 2022) and vital rates (Pletscher et al., 1997, Ausband & Mitchell, 2021) as a population transitions from recolonizing to established. Because our data come from a recolonizing population, our predicted dispersal probabilities and vital rates may be higher than those from established populations, leading to more optimistic population projections. However, we also may have underestimated reproduction, as our average number of pups (1.50 six-month-old pups/pack) is reflective of smaller pack sizes and we did not allow for multiple breeding events, which occur in larger wolf packs (Ausband, 2018). We did not have sufficient data to detect sex-specific effects in our estimation model. As a result, we did not model the sex of individuals, and we simply assumed that a pack forms when there are ≥2 adults in a territory. However, previous work suggests that established wolf populations may have slightly male-biased sex ratios (Ausband, 2022) and dispersal (Ausband, 2022; Jimenez et al., 2017). The additional data to support additional complexity should come with ongoing monitoring of this population.

Our spatially explicit projection model fills an important but previously underexplored niche in population modeling, in which spatially explicit projections are of interest but there is insufficient information to develop a fully spatial IPM (sensu Chandler and Clark 2014). The flexibility of our modeling approach allows our framework to be applied given different types of data on demography and movement (e.g., tracking data, mark-recapture data), different types of movement models (e.g., movement in pairs or groups; Scharf & Buderman, 2020), and different approaches to modeling the suitability (e.g., dynamic ensemble models; Abrahms et al. 2019) or connectivity (e.g., resistance kernel; Kaszta et al. 2019) of habitat.

Non-stationary populations are the rule rather than the exception. Populations that are changing in distribution are of growing interest to ecologists and managers, as predators reclaim areas from which they have been extirpated (e.g., Chapron et al. 2014), as invasive populations continue to spread (e.g., Crowl et al. 2008), and as a variety of species exhibit climate-induced range shifts (e.g., Chen et al. 2011). We anticipate growing interest in the development of integrated estimation and projection approaches that accommodate significant spatial flux in populations (e.g., Elith et al. 2010), as managers are tasked with predicting such shifts and responding to the management challenges they may pose (e.g., Synes et al. 2015).

## Supporting information

Petracca et al. Appendix 1

Petracca et al. Appendix 2

Petracca et al. Appendix 3

Petracca et al. Appendix 4

Petracca et al. Appendix 5

## ACKNOWLEDGMENTS

We thank the Washington Department of Fish and Wildlife (WDFW) for providing funding for this work, and the Washington Cooperative Fish and Wildlife Research Unit for facilitating the funding. We deeply appreciate the insights and advice of WDFW scientists and managers, particularly D Martorello, T Roussin, J Smith, and G Spence, and we thank the members of the Washington Fish and Wildlife Commission, particularly the Wolf Committee, for challenging us to continually improve this product. Members of the Quantitative Conservation Lab and the Quantitative Ecology Lab at the University of Washington provided advice and support throughout. Any use of trade, firm, or product names is for descriptive purposes only and does not imply endorsement by the U.S. Government.

