## Supplementary material for "Merging integrated population models and individual-based models to project population dynamics of recolonizing species": Petracca et al. Appendix 1

**Appendix S1. Details on trapping and surveying of wolves in Washington, USA**

Upon capture, demographic and morphometric data were collected, and wolves were fitted with GPS- and radio telemetry-equipped collars programmed to collect two to six GPS fixes per day. Wolves were collared in two seasons, the aerial capture season (mid-January through early March) and the summer trapping season (May to early September). Aerial captures targeted packs with collars already present by using radio-telemetry signals to help locate packs, after which animals were darted from helicopters. Summer captures targeted packs without collars, with wolf presence suspected due to physical sign (e.g., tracks, dens) or reports from the public. For summer captures, WDFW biologists placed two to five camera traps within a 350-km^2^ area over a 9- to 14-day period to ascertain wolf presence. If wolves were confirmed present, biologists used trap lines with traps located every ~0.8 km along putative wolf travel routes, with baited scented lures, to capture animals, which were then fitted with collars. See WDFW (2019) for details on wolf trapping.

Aerial surveys targeted packs with at least one collared individual, with biologists counting all observed wolves from helicopters or fixed-wing aircraft. From 2009-2014, biologists also noted the number of six-month-old pups in each pack during aerial surveys. Track surveys and camera surveys targeted suspected packs without collared individuals. Track surveys were 20 to 110 kilometers in length, relying on detection of wolf tracks on suitable substrate (i.e., dirt, snow) to confirm wolf presence and number of individuals. Camera surveys were conducted by placing cameras along roads or trails that served as common travel routes for wolves.
