## Supplementary material for "Merging integrated population models and individual-based models to project population dynamics of recolonizing species": Petracca et al. Appendix 2

**Appendix S2. Identification of wolf movement states from telemetry data**

***Identification of movement states***

We used package adehabitatLT (Calenge, 2015a) in program R (R Core Team 2022) to rarefy GPS collar data to daily locations closest to 07:00 local time each day, a time when wolves were assumed to be most active (Theuerkauf et al. 2003). We started a new trajectory when there was a re-collaring event of an individual, and removed trajectories with <60 days of data. We had a total of 31,607 daily points across 83 movement trajectories from 74 wolves.

Based on the GPS collar data, we calculated first passage time (FPT) and net squared displacement (NSD) to identify movement states of individuals. Based on these metrics, we delineated segments of each individual’s complete movement trajectory during which the individual was a “mover” or a “resident.” Broadly, a “mover” is an animal that is moving relatively long distances quickly, while a “resident” is an animal that is maintaining a territory.

For each of 83 trajectories, we used package adehabitatLT (Calenge 2015a) to regularize the trajectories, a necessary step to calculate FPT, the time required for an animal to cross a circle of radius *r* (Johnson et al. 1992). Radius *r* was calculated for each individual as the scale in the movement data with maximum variance of log(FPT). This dominant scale of variability indicates the scale at which behaviors are occurring at the home range level, such as area-restricted search behavior (Fauchald and Tveraa 2003). We identified this radius for each individual using the fpt and varlogfpt functions in package adehabitatLT (Calenge 2015a), and then calculated the median radius across individuals. We then used this radius (18 km) to calculate FPT along the trajectory of telemetry locations for each collared wolf. Five trajectories (6% of total) could not be analyzed via FPT at that scale because movements were too localized, leading to a classification of these as resident trajectories.

We used the penalized contrast method of Lavielle (1999, 2005) to perform a non-parametric segmentation of each movement trajectory, using FPT at an 18-km radius as our path signal. We used the lavielle function in package adehabitatLT (Calenge 2015a) to compute the matrix used to segment the series, where the optimal number of segments was the segment value representing 80% of the range of the contrast function. The limits of each segment were determined using the findpath function of package adehabitatLT (Calenge 2015a).

After segmentation, we calculated for each segment [1] maximum NSD and [2] median FPT. We determined that segments where NSD was >75th percentile and FPT was >50th percentile best discriminated movers. When compared to assessments by the Washington Department of Fish and Wildlife Wolf Specialist (BTM), these trajectories represented dispersal (i.e., permanent movement from an origin territory to a new area; Gese and Mech 1991, Jimenez et al. 2017) 84.6% of the time (6.4% omissions, 9.0% commissions). Omissions were due to shorter dispersal events that were imperceptible using NSD, and commissions occurred due to large exploratory movements that did not terminate in a new territory. All movement segments were removed from wolf trajectories for calculation of home range and resource selection but were retained for defining the state of an individual in our multi-state survival model.

***Identification of dispersal distance and territory size***

We calculated movement distances of movers as the Euclidean distance between the first and last points of each segment in which a given wolf was a mover. We modeled these distances with a gamma distribution and sampled from the resulting estimates to determine the distances moved by potential dispersers in the projection model.

We also used daily GPS collar data, excluding “mover” segments as described above, to calculate mean territory size of wolves in Washington. These data included 23,965 daily points from 74 wolves. We first used the package adehabitatHR (Calenge 2015b) to calculate 99% minimum convex polygons (MCPs) for individual wolves between May 1 and April 30 of the following year. We used the 99% MCP approach to be liberal in identifying the area defended by wolves. We subset the MCPs further to only include those with greater than 50% of their area in Washington, and then combined MCPs from individuals in the same pack in the same year. For analysis of mean territory size, we included only those MCPs with a minimum number of 180 points, as this minimum value removed a positive association between number of points and annual home range size (𝛽 = 0.33, SE = 0.38, *p* = 0.393 at $n\geq180$). We calculated mean territory size using a linear mixed model in package lme4 (Bates et al. 2015), with pack as a random effect.
