## Supplementary material for "Merging integrated population models and individual-based models to project population dynamics of recolonizing species": Petracca et al. Appendix 3

**Appendix S3. Description of survival, abundance, reproduction, and population process models used in the Integrated Population Model (IPM) to estimate wolf population dynamics in Washington, USA**

*Survival and state transitions*

We used a known-fate multi-state model with staggered entry and a six-month time step (Brownie et al., 1993; Kaplan & Meier, 1958; Pollock et al., 1989) to estimate survival and state transitions. The four possible states were [1] alive and resident within Washington, [2] alive and moving within Washington, [3] alive and moving outside of Washington, and [4] dead/permanently emigrated. Mortalities due to removals by the state agency in response to livestock depredations (hereafter “removals”) were censored rather than recorded as mortalities, which allowed us to directly adjust management removals in population projections. Resident (i.e., non-mover) wolves that resided in territories located on the border of Washington and another state or province sometimes left Washington, but were considered to be in state 1. A wolf was declared to be in one of the movement states (States 2 or 3) if it was classified as a mover at any point within a given six-month interval, based on the analysis of its segmented movement trajectory as described in Appendix S2. To enter state 3, animals had to leave Washington during an interval in which they were classified as a mover. In our dataset, only one collared animal that left Washington while a mover returned, and because of the rarity of this event, we forced individuals in state 3 to transition deterministically to state 4, the absorbing state.

The observation data were modeled as:

${obs}_{i,t} \sim Categorical(\gamma_{g_{i,t-1,}{obs}_{i,t-1},1:4,t})$,	(Eq. 1)

where ${obs}_{i,t}$ represents the observed state of individual $i$ at occasion $t$, and $\gamma$ is the transition array:

|  |  | *t* + 1 | | | |
| --- | --- | --- | --- | --- | --- |
|  |  | Resident | Moving, in WA | Moving, not in WA | Dead |
| *t* | Resident | $\varphi_{g,t}^{A}*\left( 1- \epsilon_{g}^{A} \right)$ | $\varphi_{g,t}^{A}* \epsilon_{g}^{A}* \alpha$ | $\varphi_{g,t}^{A}* \epsilon_{g}^{A}*(1- \alpha)$ | $1- \varphi_{g,t}^{A}$ |
|  | Moving, in WA | $\varphi_{g,t}^{B}*\left( 1- \epsilon_{g}^{B} \right)$ | $\varphi_{g,t}^{B}* \epsilon_{g}^{B}* \alpha$ | $\varphi_{g,t}^{B}* \epsilon_{g}^{B}*(1- \alpha)$ | $1- \varphi_{g,t}^{B}$ |
|  | Moving, not in WA | 0 | 0 | 0 | 1 |
|  | Dead | 0 | 0 | 0 | 1 |

where $\varphi_{g,t}^{A}$ is survival of a resident, $\varphi_{g,t}^{B}$is survival of a mover, $\epsilon^{A}$is the probability of initiating movement*,* $\epsilon^{B}$ is the probability of continuing movement, and $\alpha$ is the probability of a moving wolf staying in Washington. Wolves were aged in the model in six-month increments, but multiple ages were lumped into age groupings ($g$) for purposes of demographic rate estimation, given sample size limitations. In particular, $\varphi_{g=juv}$ is the probability of surviving a six-month interval for animals between 6 and 24 months of age and $\varphi_{g=adult}$ is the probability of surviving a six-month interval for animals that are 24+ months old. The same age groupings were used for $\epsilon_{g}^{A}$ and $\epsilon_{g}^{B}$, except that wolves were not permitted to enter a movement state before 12 months of age, and thus $\epsilon_{g}^{A}$ was set to zero for animals <12 months old.

Few data were available to inform $\epsilon^{B}$(only six out of 83 mover trajectories exhibited a termination of movement while in Washington), which necessitated use of a more informative prior for this parameter. Mean duration of dispersal events reported in Jimenez et al. (2017) was 5.5 months (SD = 6.8); we used this information to create an informative prior of $Beta$(36,84) for $\epsilon^{B},$ which resulted in a similar distribution of dispersal times in our six-month-interval model (see below). The parameters $\alpha$ and $\epsilon^{A}$ were given vague priors of $Beta$(1,1).

We modeled survival, $\varphi_{g,t}^{K}$, for animals in state *K* as:

$$logit\left( \varphi_{g,t}^{K} \right)= \mu_{g}^{K}+ ϛ_{t}$$

$$ϛ_{t} \sim Normal\left( 0,\sigma_{period} \right),$$

(Eq. 2)

where $ϛ_{t}$was the common, across state *K* and age grouping $g$, random effect of six-month period *t* with standard deviation $\sigma_{period}$. Both $\mu$ and $\sigma_{period}$ were given vague priors$: \mu_{g}^{K}$ ~ $Normal$(0, 10) and $\sigma_{period}$~ $Uniform$(0,10).

*Development of an informative prior for the probability of continuing in the movement state,* $\epsilon^{B}$

Based on 182 wolves with known starting and ending dates for their dispersal events, Jimenez et al. (2017) reported a mean dispersal duration of 5.5 months with SD = 6.8 months and a range of 1 day – 26 months. We simulated movement data for 182 individuals on a monthly scale. Individuals started as “movers” in month 1 and then remain moving the next month with probability, $\sqrt[6]{\epsilon^{B}}$ (converting the 6-month probability from our model to a 1-month probability). We simulated individual wolf movement states for 50 months, once a wolf stopped moving, they were not allowed to start moving again in the simulation.

For $\epsilon^{B}$, we selected a random value based on a Beta distribution, and based on our model description, we held $\epsilon^{B}$ constant for each month and across wolves within a simulation scenario. We used a grid search-based approach, simulating the process above for all combinations of values of alpha and beta from 1 to 100. For each combination of alpha and beta, we simulated 1000 datasets and summarized the results as a mean, standard deviation, and maximum number of months in a movement state.

The results were very similar for many of the parameter combinations that resulted in mean of the Beta distribution that was ~0.3; all combinations that were close to the mean (5.5) underestimated the standard deviation (~5 months in the simulation) and overestimated the maximum (~29 months) when compared to Jimenez et al. (2017). To compare the monthly results with our six-month time frame, we also simulated movement state data under a six-month interval and found the results were very similar. Due to the similarity of results, we used the values for the Beta distribution that were closest to the mean of 5.5 in the monthly simulation.

*Abundance model*

We modeled pack count data to estimate abundance, ${Ntot}_{t,s}$, at time $t$ in territory *s* as:

$\log\left( C_{t,s}+1 \right)\sim Normal(\log\left( {Ntot}_{t,s}+1 \right), \sigma_{pack}$),	(Eq. 3)

,

where $C_{t,s}$ was the end-of-year minimum counts of wolves observed by WDFW at time $t$ and territory $s$. We put an $Inverse Gamma$(1,1) prior on the variance of counts, ${\sigma_{pack}^{2}}$. With this prior, a miscount of one wolf was most probable and miscounts of 4 or fewer wolves represented ~80% of the prior distribution. Note that because pack and pup counts occurred once per year in December, we only fit this model and the reproduction model below when $t=1,3,5,\ldots.23$.

*Reproduction model*

The number of six-month-old pups in December of each year (for $t=1,3,5,\ldots.23$) at territory $s$ ($R_{t,s})$ was modeled as

$$R_{t,s} \sim Categorical\left( \pi_{1:7} \right)-1 if \geq2 adults in the territory$$

$$R_{t,s}=0 if<2 adults in the territory$$

(Eq. 4)

where $R_{t,s}$ was NA (and therefore estimated by the model) if pup counts were not recorded but >= 2 adults were in that territory, and $\pi_{1:7}$ was the vector of probabilities that a wolf territory had from 0-6 six-month-old pups. We gave $\pi_{1:7}$a vague Dirichlet prior reflecting equal probability of 0-6 pups.

*Population process model*

In each territory $s$ that was occupied during the data collection period, we modeled age-specific wolf abundance beginning in the year wolves were first counted in that territory. The first sampling period in the population process model was December 2009, and new territories could enter the model in December of subsequent years (i.e., $first_{s}\in1,3,5,\ldots.23)$.

Total pack size in the first year that a pack occupied a given territory $s$ was modeled as:

${Ntot}_{first_{s},s} \sim Poisson(N_{init})$	(Eq. 5)

where $N_{init}$ had a $Uniform$(0,20) prior. The number of individuals in each age class in the first time period followed a multinomial distribution such that,

$N_{1:3,first_{s},s} \sim Multinomial\left( {SA}_{1:3}\boldsymbol{,}{Ntot}_{first_{s},s} \right)$	(Eq. 6)

where ${SA}_{1:3}$ = {0.123, 0.114, 0.763} was the stable age distribution, calculated as the scaled dominant eigenvector of a Leslie matrix. Fecundity in the Leslie matrix was estimated from the observed pack and pup count data and represented the median number of six-month-old pups per older wolf. Annual survival probabilities in the Leslie matrix were estimated from an independent run of the multi-state model described above. For $t > first_{s}$, the total number of wolves in six-month period *t* in territory *s* was calculated as the sum of the number of individuals in each age class:

${Ntot}_{t,s}= N_{1,t,s}+ N_{2,t,s}+ N_{3,t,s}.$	(Eq. 7)

Due to seasonal differences in how data were collected and how animals aged in the model, we had different process models for odd numbered periods (i.e., December) and even numbered periods (i.e., June). For $t=1,3,5,\ldots.23$, the abundance of wolves in the first age class was calculated as:

$N_{1,t,s}={\lambda.pups}_{t,s}- {removals}_{1,t,s}$	(Eq. 8)

where ${removals}_{1,t,s}$ are animals that were removed by the state agency prior to 6 months of age. For $t=2,4 ,6\ldots22$, the abundance of wolves in the first age class was modeled as:

$N_{1,t,s}\sim Binomial(\varphi_{juv, t-1}^{A} *(1- \epsilon_{juv}^{A}), N_{1,t-1,s})$	(Eq. 9)

using random binomial draws to account for demographic stochasticity. Note that $g$ subscripts on $\varphi$ and $\epsilon$ refer to age groupings for the multi-state model as described above.

For all $t$, abundances for age classes 2 and 3 were calculated as:

$N_{age,t,s} = {N.residents}_{age,t,s}- {removals}_{age,t,s}+ {N.immig}_{age,t,s}+ {N.dispersers}_{age,t,s}$. (Eq. 10)

That is, these age classes were composed of residents who survived and remained ($residents$) minus individuals that were removed by the state agency $(removals)$ plus immigrants entering Washington from out of state $(immig)$ plus dispersers from within Washington who joined the pack $(dispersers)$. Removals for each age class, time period, and territory, ${removals}_{age,t,s}$, were provided as data by Washington Department of Fish and Wildlife.

For $t=1,3,5,\ldots.23$, the number of residents in age classes 2 and 3 were calculated based on survival and state transitions of animals that were previously in the pack:

${N.residents}_{2,t,s} \sim Binomial(\varphi_{juv,t-1}^{A}*(1- \epsilon_{juv}^{A})$ , $N_{1,t-1,s})$	(Eq. 11)

${N.residents}_{3,t,s} \sim Binomial(\varphi_{juv,t-1}^{A}*(1- \epsilon_{juv}^{A})$ , $N_{2,t-1,s})$ + $Binomial(\varphi_{ad,t-1}^{A}*(1- \epsilon_{ad}^{A})$ , $N_{3,t-1,s})$	(Eq. 12)

For $t=2,4 ,6\ldots22$, Eq. 11 remains the same, but Eq. 12 is modified such that:

${N.residents}_{3,t,s}\sim Binomial(\varphi_{ad,t-1}^{A}*(1- \epsilon_{ad}^{A})$ , $N_{2,t-1,s}+ N_{3,t-1,s})$	(Eq. 13)

i.e., all animals entering age class 3 in June survived with the probability for resident adults.

To estimate the number of out-of-state immigrants, we followed Abadi et al. (2010), such that for $t>first_{s}$:

${Tot.immig}_{t,s}\sim Poisson\left( \lambda_{immig} \right)$	(Eq. 14)

and

${Nimmig}_{1:3,t,s}\sim Multinomial\left( {Immig}_{1:3}, {Tot.immig}_{t,s} \right)$	(Eq. 15)

where $\lambda_{immig}$ is the average number of immigrants to each pack per time interval with prior $Uniform(0,5)$. To represent the age distribution of immigrants, we used the age distribution of 33 wolves observed in Washington that became movers, ${Immig}_{1:3}$ = {0, 0.3, 0.7}, i.e., 70% of these individuals were in age class 3.

To calculate in-state dispersers that joined territories, we first had to model the number of individuals that were in state 2, the in-state movers. To do so, at each time step we added the total number of new movers in each age class to the number of movers that continued in the movement state in each age class. For example, for age class 1 when $t > first_{s}$, the number of individuals moving at time $t$ is modeled as:

${N.movers}_{1,t}\sim Bin(\varphi_{juv,t-1}^{A}* \epsilon_{juv}^{A}* \alpha$, $N_{1, t-1,1:s})$ + $Bin(\varphi_{juv,t-1}^{B}* \epsilon_{juv}^{B}* \alpha$, ${N.movers}_{1, t-1}$)

(Eq. 16)

The model is analogous for age classes 2 and 3 where the appropriate age classes, survival probabilities, and movement probabilities are used. We note that in December of each year ($t = 1,3,5,...23$), age classes 2 and 3 will also include individuals from the previous age class that aged into that class.

The number of potential dispersers was then modeled based on the movers that terminated movement:

${N.potential.dispersers}_{age, t} \sim Binomial(\varphi_{g,t-1}^{B}$ $*\left( 1- \epsilon_{g}^{B} \right)$, ${N.movers}_{age,t-1})$	(Eq. 17)

From this, we modeled the number of dispersers, by age, joining any territory, $s$:

${N.dispersers}_{age,t,s} \sim Multinomial({settle}_{t,s}$, ${N.potential.dispersers}_{age,t})$	(Eq. 18)

where $settle$*,* the probability of settling in territory $s$, was equal across all territories occupied at time $t$. This equation captures a primary difference between the model in the data collection period and the model in the projection period. In the data collection period, potential dispersers could not settle in unoccupied territories and the colonization process of in-state dispersers was entirely latent. By contrast, in the projection period, the colonization process included unoccupied territories and was modeled explicitly (see *Individual-Based Model*, below). We further note that [1] there were no potential dispersers in June of the first sampling year from a given territory, as wolves cannot start moving until this period and potential dispersers are required to have moved for at least one six-month period, and [2] there were no potential dispersers aged up to and including 12 months in our model, as wolves could only start moving at 12 months and are ineligible to settle until they are 18 months.
