## Supplementary material for "Merging integrated population models and individual-based models to project population dynamics of recolonizing species": Petracca et al. Appendix 4

**Appendix S4. Habitat suitability of potential wolf territories via second-order resource selection function (RSF) and Least Cost Path computation**

We used a resource selection function (RSF) of the second order (Manly et al. 2002) to determine the relative suitability of wolf territories for wolf colonization in Washington. Our “used” surface comprised 17,907 daily locations of wolves within 112 99% minimum convex polygons (MCPs) of annual pack territories. These territories had a minimum of 60 daily GPS points and greater than 50% of their area within Washington. Our “availability” surface included both the 112 MCPs and all land area within a radius of 31.11 km (the diameter of an average-sized (760 km^2^) wolf territory in Washington) from each MCP (Figure S1). We sampled availability to used points in a 20:1 ratio, ensuring that this ratio was maintained within each MCP. In total, there were 358,140 available points and 17,907 used points in the analysis, for a total of 376,047 points.

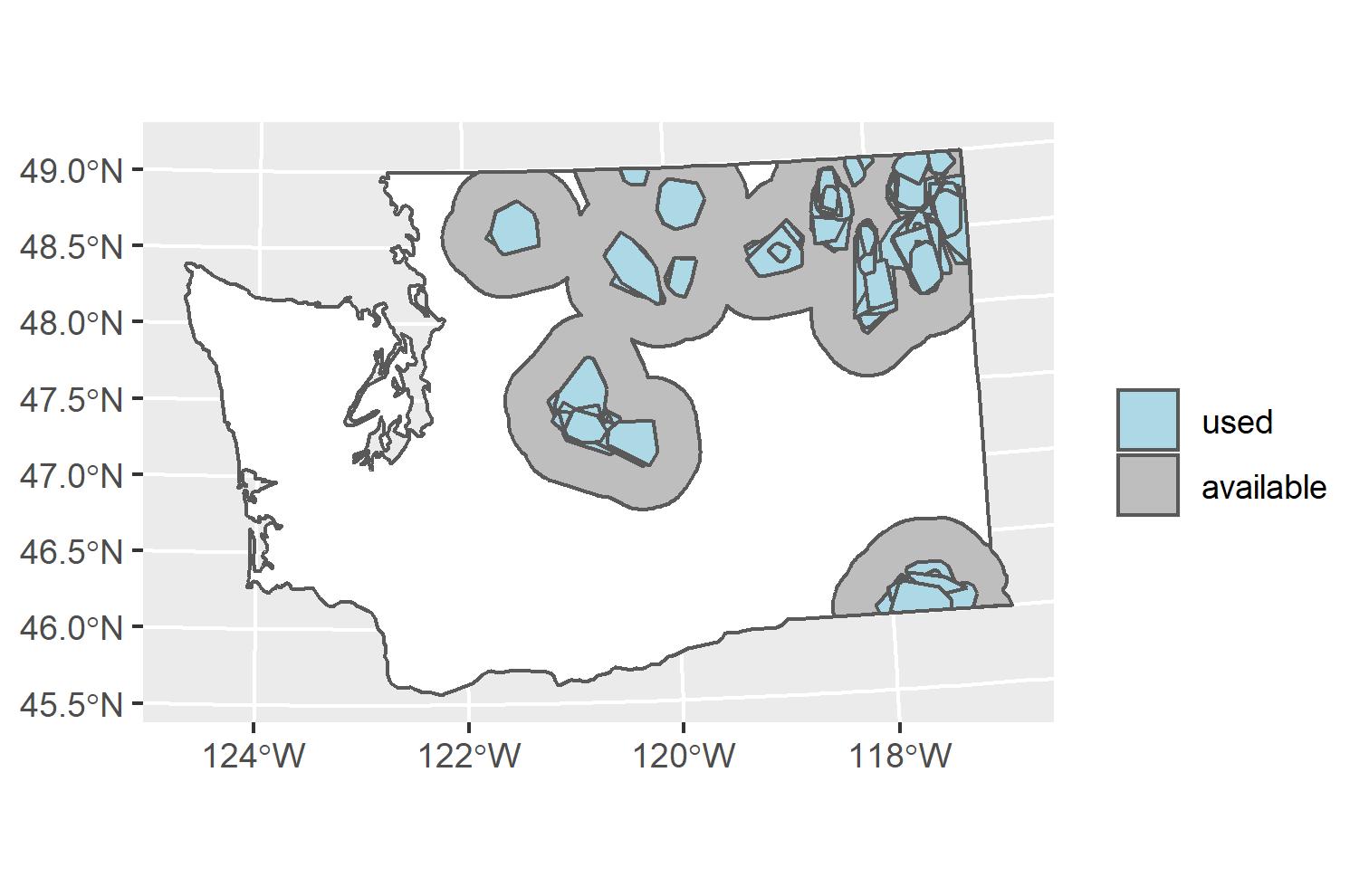

Figure S1. Summary of used and available area for the second-order resource selection function (RSF) for wolves in Washington State, USA, based on daily GPS collar data from 74 individuals.

Covariates of interest were human population density (2010 TIGER data; U.S. Census Bureau), agricultural cover (0/1) (NLCD 2016; CONUS), grassland cover (0/1) (NLCD 2016; CONUS), forest cover (0/1) (NLCD 2016; CONUS), shrubland cover (0/1) (NLCD 2016; CONUS), terrain ruggedness (Welty & Jeffries 2018)(Brooks et al. 2017), road density (meters of road per km^2^; WSDOT), elevation (30-m SRTM GL1; NGA and NASA), relative deer abundance (WDFW), distance to state highway (WSDOT), and presence of public grazing allotment (0/1) (WDFW; BLM; USFS). Relative deer abundance corresponded to the number of white-tailed deer, black-tailed deer, and mule deer harvested within each Game Management Unit (GMU) in Washington from 2017-2019, with the “Close Gaps” spatial interpolation method in SAGA GIS (Conrad et al. 2015) used to infer relative deer abundance in non-GMU areas. All rasters were standardized to a resolution of 30 meters. We used the extract function in R package raster (Hijmans & van Etten 2014) to sample covariate values at each used and available point.

The RSF was run using R package glmmTMB (Brooks et al. 2017) with a logistic link, pack-specific intercepts with large, fixed prior variance (${\sigma_{\alpha}}^{2} = 10^{6})$, and weight assignment of 100 to available points (Muff et al. 2020). All covariates were standardized prior to inclusion in the model. Linear relationships were predicted for all covariates but elevation and terrain ruggedness, which incorporated a quadratic effect. In addition, distance to highway was modeled with a decay function (e.g., log(x)+1; see Prokopenko et al. (2016)), such that the impacts of roads would lessen as distance increased. The global model for $\pi_{nj}$, the probability that a point $y$ will be used by pack $n$ at location *j* was,

logit($\pi_{n,j})$= ${(\beta}_{1}$ * ${agriculture}_{n,j})$ + ${(\beta}_{2}$ * ${allotment}_{n,j})$ + ${(\beta}_{3}$ * ${deer}_{n,j})$ + ${(\beta}_{4}$ * ${\log\left( disthwy+1 \right)}_{n,j})$ + ${(\beta}_{5}$ * ${elevation}_{n,j})$ + ${(\beta}_{6}$ * ${{elevation}_{n,j}}^{2})$ + ${(\beta}_{7}$ * ${forest}_{n,j})$ + ${(\beta}_{8}$ * ${grassland}_{n,j})$ + ${(\beta}_{9}$ * ${humanpop}_{n,j})$ + ${(\beta}_{10}$ * ${roaddensity}_{n,j})$ + ${(\beta}_{11}$ * ${ruggedness}_{n,j})$ + ${(\beta}_{12}$ * ${{ruggedness}_{n,j}}^{2}$) + ${(\beta}_{13}$ * ${shrubland}_{n,j})$+ $\alpha_{n,}$

where $\alpha_{n}$ was a pack-specific intercept for pack $n$.

Wolves were more likely to select home ranges with greater relative deer abundance (𝛽 = 0.32, SE = 0.01), forest cover (𝛽 = 0.10, SE = 0.04), shrubland cover (𝛽 = 0.11, SE = 0.02), distance from state highway (𝛽 = 0.39, SE = 0.01), and where public grazing allotments were present (𝛽 = 0.52, SE = 0.02) (Figure S2). Wolves were also more likely to select home ranges in areas with lower human population density (𝛽 = -1.16, SE = 0.05), agricultural cover (𝛽 = -0.49, SE = 0.03), road density (𝛽 = -0.12, SE = 0.01), grassland cover (𝛽 = -0.10, SE = 0.02), and terrain ruggedness (𝛽 = -0.46, SE = 0.01 for linear and 𝛽 = 0.05, SE = 0.01 for quadratic term). Wolves also selected for areas at intermediate elevation (𝛽 =0.27, SE = 0.02 for linear and 𝛽 = -0.47, SE = 0.01 for quadratic term) (Figure S2).

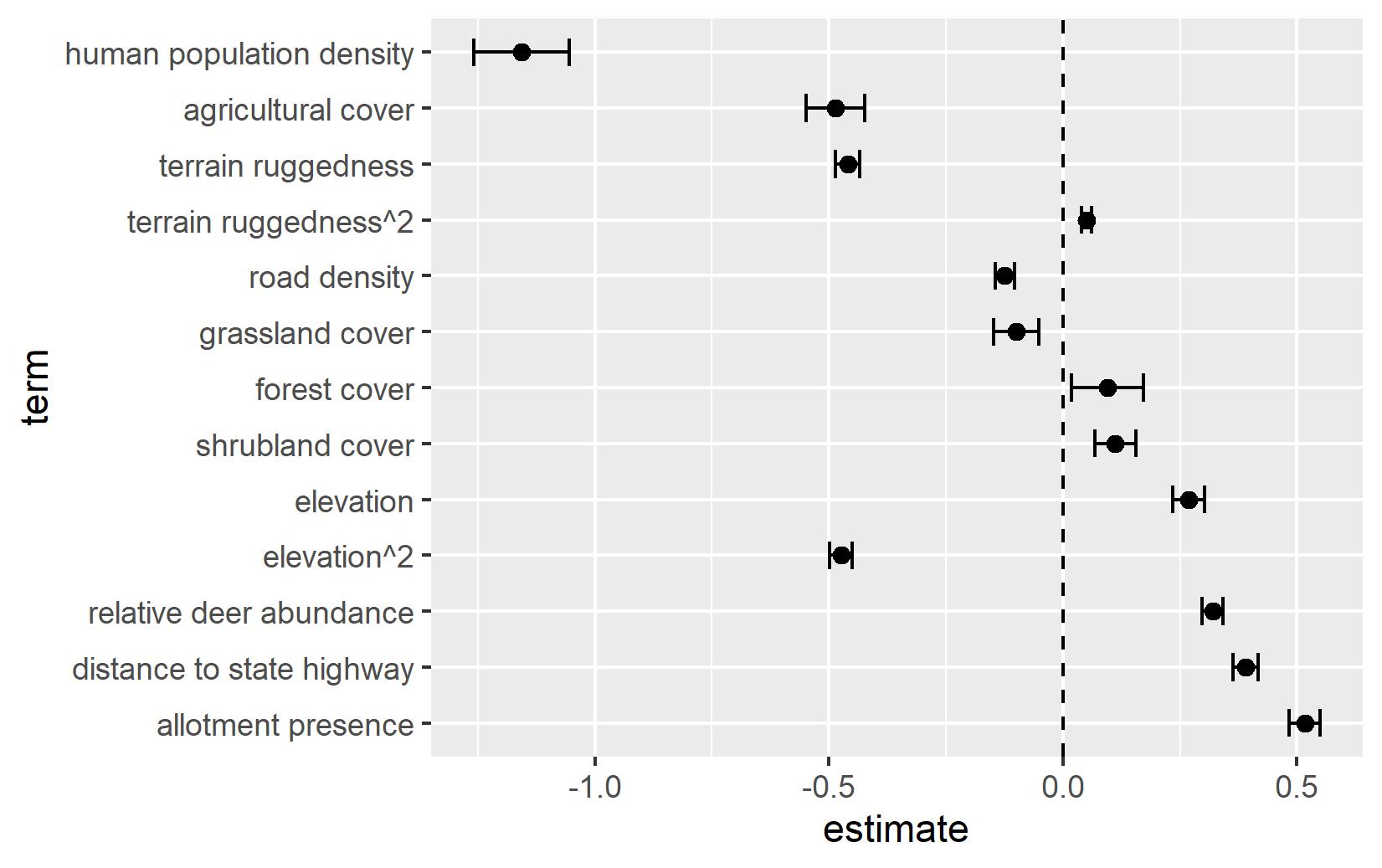

Figure S2. Beta coefficients and 95% CI for variables included in the second-order resource selection function (RSF) for wolves in Washington State, USA, based on daily GPS collar data from 74 individuals.

These covariate relationships were then extrapolated to all pixels within the state, using the mean and standard deviation of data points used in the RSF model to standardize the statewide covariate rasters. Areas of greatest relative selection for wolf territories largely followed forested, undeveloped areas, including the national forests in the Northeast, the Northern and Southern Cascades, and the Olympic Peninsula (Figure S3).

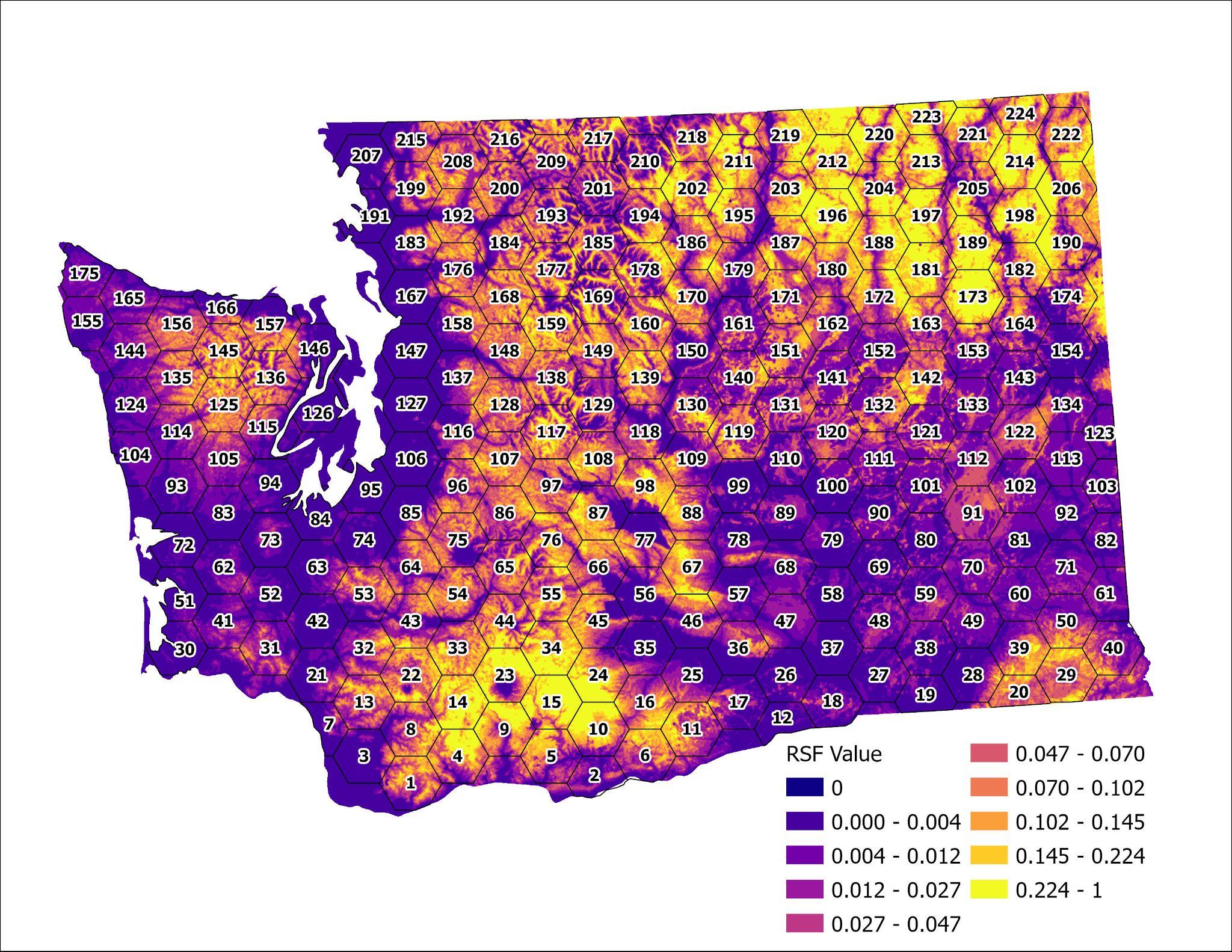

Figure S3. Predicted estimate of second-order resource selection function (RSF) for wolves in Washington State, USA, based on daily GPS collar data from 74 individuals.

The least cost path analysis was performed in Python-based program UNICOR (Landguth et al. 2012). UNICOR applies Dijkstra’s shortest path algorithm to individual-based simulations, identifying the single shortest path from every defined species location on a landscape to every other defined species location (Landguth et al. 2012). The program requires two inputs: [1] a resistance surface, and [2] specified species locations. Our resistance surface was the inverse of the RSF presented in Figure S3, where pixels of high relative selection are of low resistance to movement, and pixels of low relative selection are of high resistance to movement. Specified species locations corresponded to the centroid of each of the 224 potential pack territories in Washington.

The output of interest from UNICOR was the cost distance matrix, a 224 x 224 matrix specifying the cost distance from each territory to all others. This matrix was brought into program R (R Core Team 2022) and used to determine, for each 1-km distance from 1 km to 632 km (the maximum observed dispersal distance of wolves in Washington), the territory of least cost from each source territory. The final output was a 224 x 632 matrix consisting of the least-cost destination territory for each of 224 source territories and 632 dispersal distances. This matrix was used in the individual-based model’s movement function to identify the territory of least cost from a source territory given a land-based movement distance.
