## Supplementary material for "Merging integrated population models and individual-based models to project population dynamics of recolonizing species": Petracca et al. Appendix 5

**Appendix S5. Comparison of model output from Categorical Resource Selection Function (RSF) and Least Cost Path territory selection processes**

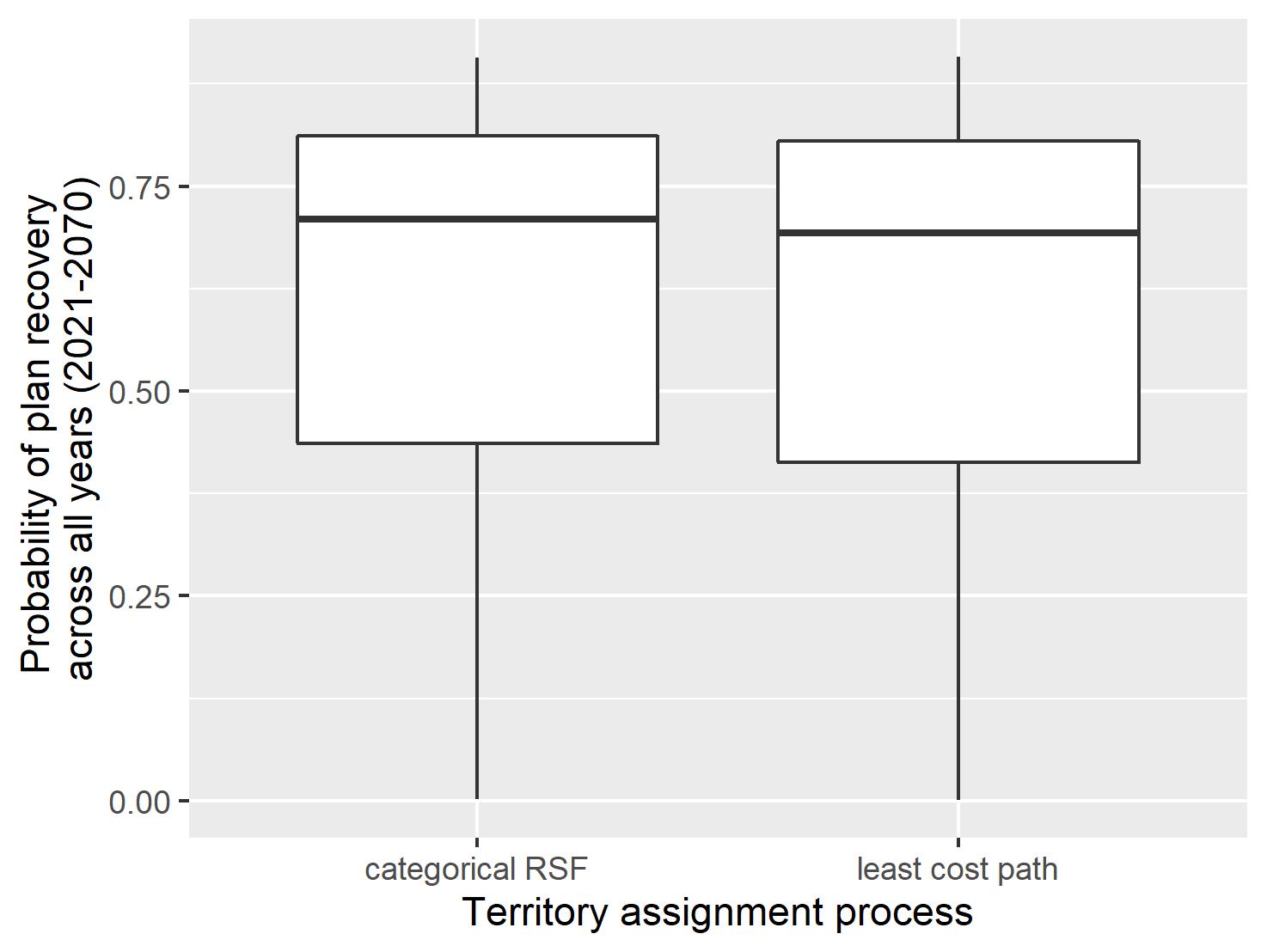

Figure S1. Probability of meeting plan recovery (i.e., having four breeding pairs in each of three recovery regions, and 15 breeding pairs total in the state) across all years (2021-2070) for two territory selection processes for wolves in Washington State, USA: a categorical resource selection function (RSF) process (left) and least cost path process (right).

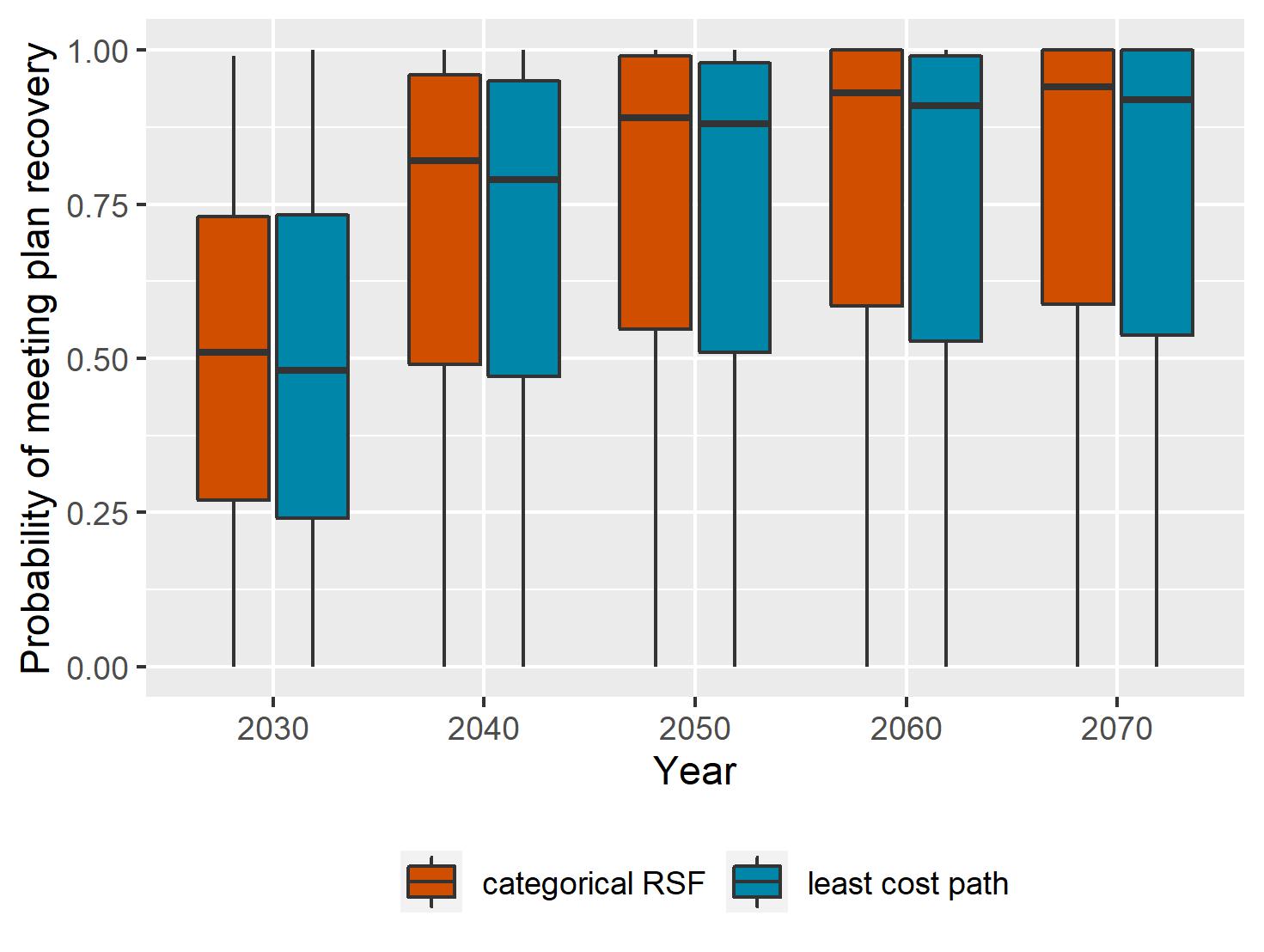

Figure S2. Probability of meeting plan recovery (i.e., having four breeding pairs in each of three recovery regions, and 15 breeding pairs total in the state) at five time steps for two territory selection processes for wolves in Washington State, USA: a categorical resource selection function (RSF) process (red) and least cost path process (blue).

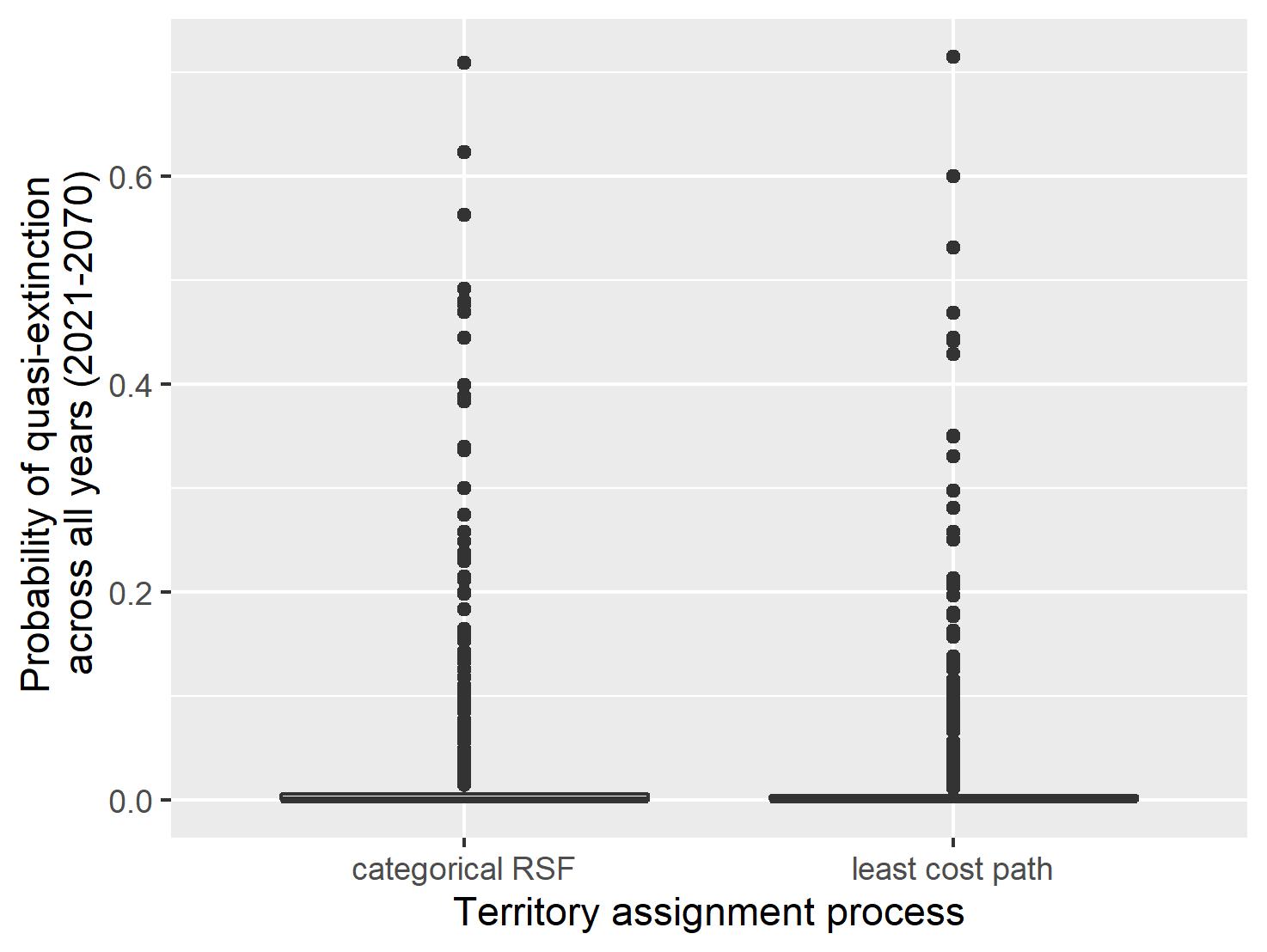

Figure S3. Probability of quasi-extinction (i.e., having <24 breeding adults in each of three recovery regions, and <92 breeding adults total in the state) across all years (2021-2070) for two territory selection processes of interest for wolves in Washington State, USA: a categorical resource selection function (RSF) process (left) and least cost path process (right). A note that the black lines at 0.0 include both the median and 50% prediction interval, and that the black lines represent outlier points from 50,000 samples.

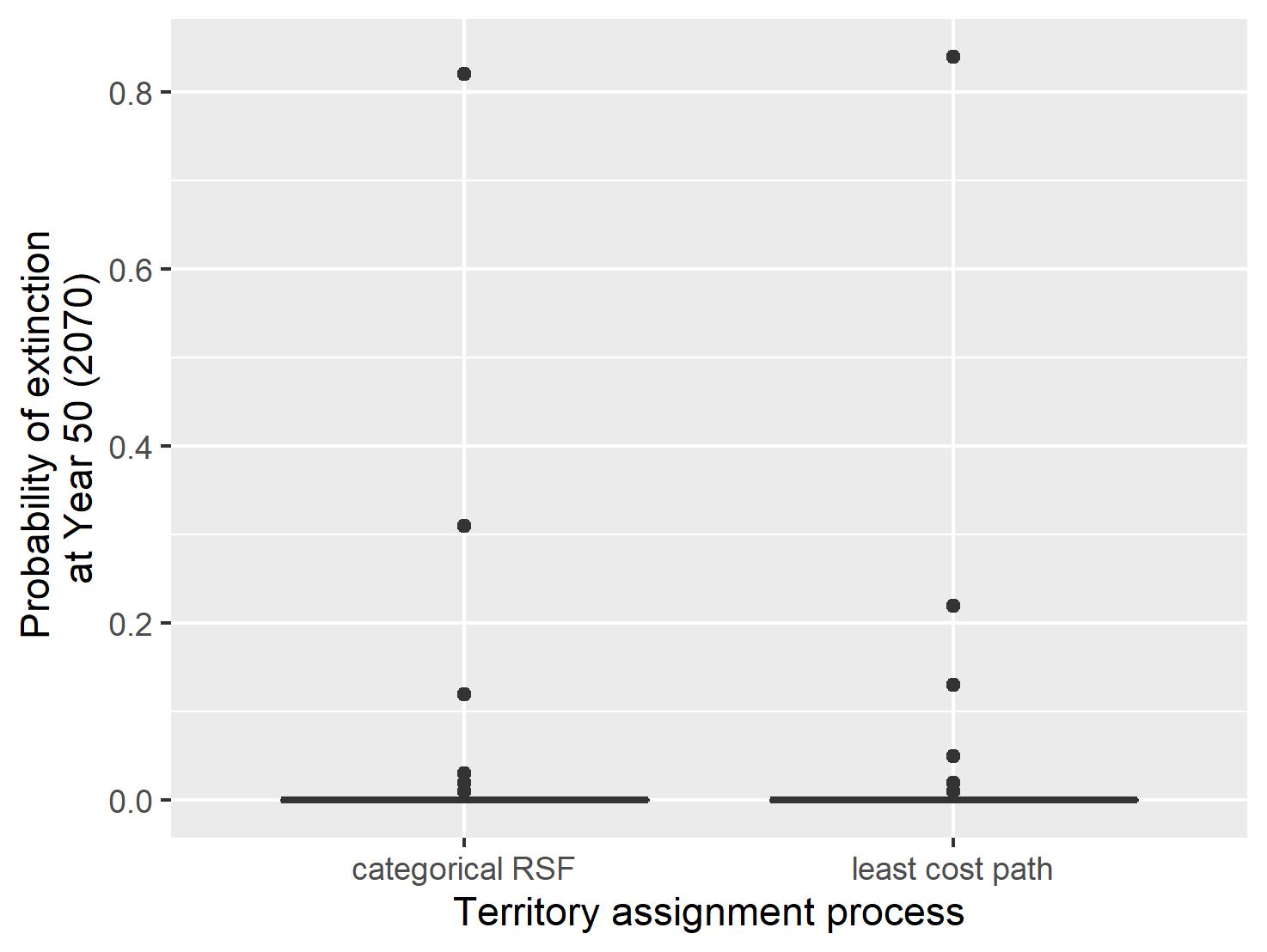

Figure S4. Probability of extinction at year 50 (i.e., having 0 wolves in year 2070) for two territory selection processes for wolves in Washington State, USA: a categorical resource selection function (RSF) process (left) and least cost path process (right). A note that the black lines at 0.0 include both the median and 50% prediction interval, and that the black lines represent outlier points from 50,000 samples.

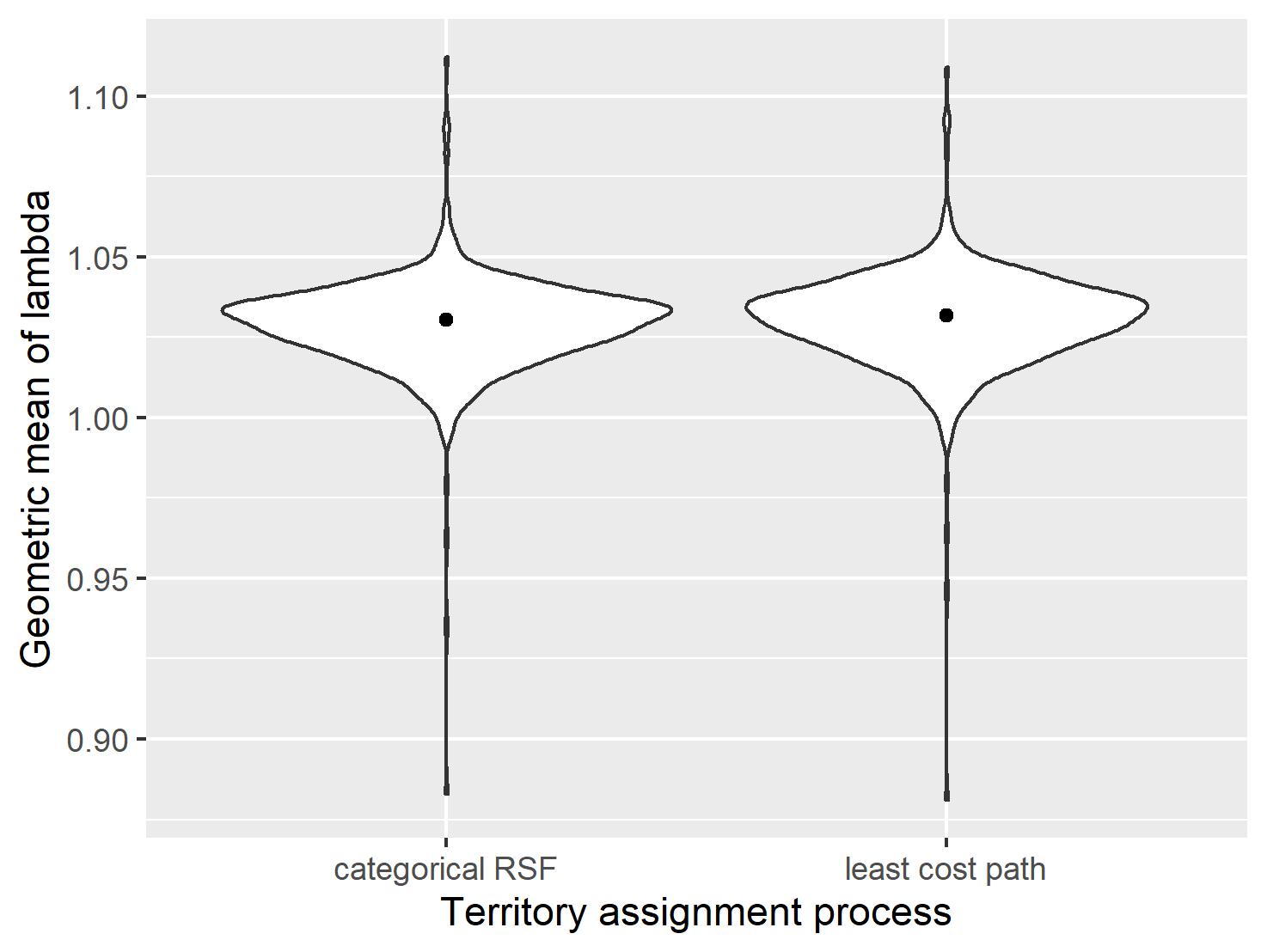

Figure S5. Geometric mean of lambda over the projection period (2021-2070) for two territory selection processes for wolves in Washington State, USA: a categorical resource selection function (RSF) process (left) and least cost path process (right).
